# MAPLE: A hierarchical Multimodal Assistant for Pathology vision-Language Evaluation using logic and reasoning

**DOI:** 10.64898/2026.11.21.768223

**Authors:** Saurabh Joshi, Jieneng Chen, Ruisheng Yuan, Varvara Wyart, Saud Al Faisal, Ranjan Mukherjee, Florin A. Selaru, Yu Shen, André Forjaz, Thom D. Greene, Donald Kramer, Eban A. Hanna, Chanhong Min, Vasco Queiroga, Kai Lewis, Victoria D. Alvarado, David N. Sokolov, Isabella Weiner, Fan Wu, Angelica Fagan, Valentina Matos-Romero, Lucie Dequiedt, Praful R. Nair, Rofieda Alwaqfi, Ralph H. Hruban, Anirban Maitra, Ashley L. Kiemen, Arrate Munoz-Barrutia, Pei-Hsun Wu, Alan Yuille, Denis Wirtz

## Abstract

Pathological diagnosis requires integration of cellular morphology, tissue architecture, and clinical context across gigapixel whole-slide images (WSIs), yet existing vision-language models remain constrained to patch-based analysis with resource-intensive architectures unsuitable for routine clinical deployment. Here we introduce MAPLE, a hierarchical pathology vision-language model that performs whole-slide diagnosis and report generation through efficient multi-scale reasoning. MAPLE employs AgenTile, a small 10-million-parameter vision transformer for cellular-level triage at 40x resolution, achieving segmentation performance equivalent to ∼65-fold larger models (e.g., CellViT++) while providing ∼95-fold faster inference. Cross-magnification integration with a CLIP-based image encoder at 10x and 5x enables recursive selection of the most diagnostically informative regions per slide. On the PathMMU benchmark (7,691 image-question pairs), MAPLE’s multimodal (image and text) reasoning module is the first to exceed pathologist performance across all four diagnostic domains (PubMed, EduContent, SocialPath, and Atlas), achieving 70.6-81.2% accuracy versus 68.3-72.9% for pathologists. MAPLE demonstrates calibrated clinical reasoning, appropriately expressing uncertainty for metastatic carcinoma requiring ancillary testing while providing a confident diagnosis for benign hamartoma. These results were validated using an additional clinical cohort. Taken together, our findings establish MAPLE as a computationally efficient, clinically deployable system for AI-augmented pathology, particularly suited to resource-limited settings.

## INTRODUCTION

Pathological assessment serves as the gold standard for diagnosing a broad spectrum of diseases, most notably cancer, where accurate evaluation determines not only the presence and type of malignancy but also guides staging, prognosis, and therapeutic strategies.^1,2^ Despite its essential role, the diagnostic process remains time-consuming and open to errors when pathologists perform repetitive tasks.^3,4^ Pathologists must comply with structured reporting templates while meticulously documenting all pertinent clinical and histopathological findings, adding considerable workload in high-volume clinical settings.^5^ Complex and rare diseases pose further challenges, demanding the expertise of specialized pathologists with years of training, experts who are often scarce in rural and resource-limited areas, resulting in inequalities in diagnostic care.^6,7^ Additionally, interpretation can be non-standardized, with diagnoses varying among pathologists in ways that affect consistency and treatment decisions.^3,4^ These challenges suggest the need for innovative solutions that streamline and unify diagnoses, and provide specialist-level diagnostic support to underserved regions.^8,9^

Recent advances in vision-language models (VLMs) offer a promising solution.^10,11^ Models such as PathChat and MUSK have demonstrated how Contrastive Language-Image Pre-training (CLIP) based vision encoders, trained on large datasets of histopathology image-text pairs (e.g., 50 million patches for MUSK), can be combined with language models to perform patch-based assessments.^10,12–19^ As these systems become increasingly specialized for medical imaging, they offer avenues to improve workflow efficiency and enhance diagnostic consistency within pathology.^20–27^

However, current VLMs, such as PathChat and MUSK, remain fundamentally constrained in their ability to process whole-slide images at full resolution.^14,18,19,28^ As a result, they are limited to analyzing small image regions and miss the contextual information needed for accurate and holistic diagnosis. Slide-level models such as TITAN and ALPaCA extend vision–language modeling to whole-slide images, but important limitations remain.^29,30^ Both operate on patch-level features from a pretrained encoder and do not explicitly segment, classify, or enumerate individual cells, so cellular morphology, cell-type composition, and cell–microenvironment relationships are captured only implicitly.^29,30^ This may limit their sensitivity to phenotypes driven by subtle cellular features, and neither provides a pathology-specific mechanism for prioritizing diagnostically relevant regions before analysis. Moreover, LLM-based slide-level systems such as ALPaCA are demonstrated on datacenter-class hardware, with inference reported on a single 80 GB NVIDIA A100 GPU, a resource not typically available in smaller laboratories and rural healthcare settings, where systems are often limited to single consumer-grade GPUs.^30^ Consequently, there remains a need for pathology-specific systems that combine cellular-level triage with multiscale tissue context while remaining deployable within the constraints of clinical computing environments.

Here, we introduce MAPLE (Multimodal Assistant for Pathology vision-Language Evaluation), a pathology-specific vision-enabled small language model designed to generate diagnostically-relevant reports from whole-slide histopathology images. MAPLE employs a hierarchical two-stage architecture: AgenTile, a lightweight 10-million-parameter vision transformer, performs cellular-level triage at 40x resolution to identify abnormal regions, while a CLIP-based image encoder extracts architectural patterns at 10x and 5x magnifications. Cross-magnification integration enables recursive selection of the most diagnostically informative regions per slide, facilitating comprehensive whole-slide analysis while maintaining computational scalability suitable for deployment on a single consumer-grade GPU, without requiring high-performance computing infrastructure. We benchmark MAPLE on PathMMU, a large-scale assessment comprising over 7,000 image-question pairs across four diagnostic domains: PubMed (scientific pathology documents), EduContent (educational YouTube teaching videos), SocialPath (Twitter posts by pathology experts), and Atlas (authoritative pathology textbooks and clinical guidelines), demonstrating that MAPLE surpasses both state-of-the-art models and expert pathologists on all tasks.^31^ We further validate MAPLE’s whole-slide diagnostic accuracy and long-form diagnostic text output on a clinical breast cohort against ground truth diagnoses, annotations, and microscopic descriptions established by a board-certified specialist and demonstrate calibrated clinical reasoning through multi-turn diagnostic dialogue. These results establish MAPLE as a computationally efficient system capable of reliable integration into routine pathology workflows, with particular relevance for resource-limited settings.

## RESULTS

### MAPLE’s architecture and training dataset

MAPLE is a computationally efficient hierarchical pathology vision-language model (VLM) that converts gigapixel whole slide images (WSIs) into diagnostically relevant visual evidence for multimodal reasoning and reporting (Fig. 1a). The workflow employs a two-stage architecture with optimal patch selection to bridge cellular and tissue-level analysis. In the first stage, AgenTile, a vision transformer (ViT), performs cell segmentation and classification of high-resolution (40x) WSIs to identify abnormal versus normal cellular regions. In parallel, the pathology-trained CLIP encoder performs zero-shot tumor classification, whereby tumor probability is predicted using text-based class descriptors rather than labelled region annotations, at a downsampled scale (10x or 5x), generating a tumor probability heatmap that captures tissue-level structural features. Each WSI is tiled into 336x336-pixel candidate patches, and a patch is eligible for selection if abnormal nuclei constitute its most frequent class, at least 90% of its area comprises viable tissue, it contains a minimum of 20 nuclei, and the CLIP encoder’s zero-shot tumor probability heatmap assigns its location a score of at least 0.70. Eligible patches are ranked by the product of tumor probability and neoplastic purity, defined as the fraction of nuclei classified as abnormal, with spatial separation enforced to ensure whole-slide coverage. By default, the 20 highest-ranked patches are retained, a number empirically determined by the GPU memory constraints of loading all three model components simultaneously on a single NVIDIA RTX 3090 GPU (AgenTile, CLIP encoder, and 14-billion-parameter language model), though this number is user-configurable to accommodate available compute resources. If fewer than 20 patches meet the eligibility criteria, patches with the highest remaining cell counts are selected in descending order to complete the selection. The three retained patches with the highest tumor probability are passed to the language model together with an overview image of the section. Slides on which fewer than 1% of tissue regions reach the tumor probability threshold are assumed to lack significant neoplastic content and are represented by the patches with the greatest epithelial and stromal cellularity for downstream analysis. In the second stage, the selected patches are aggregated and encoded via the CLIP encoder, which transforms raw pixel data into tokenized representations. A multilayer perceptron (MLP) projection layer bridges these visual tokens to the language model’s embedding space, creating PathLLM, a unified vision-language framework where patient-level queries and histopathological representations are jointly processed through cross-modal reasoning to generate diagnostic outputs.

**Figure 1:**
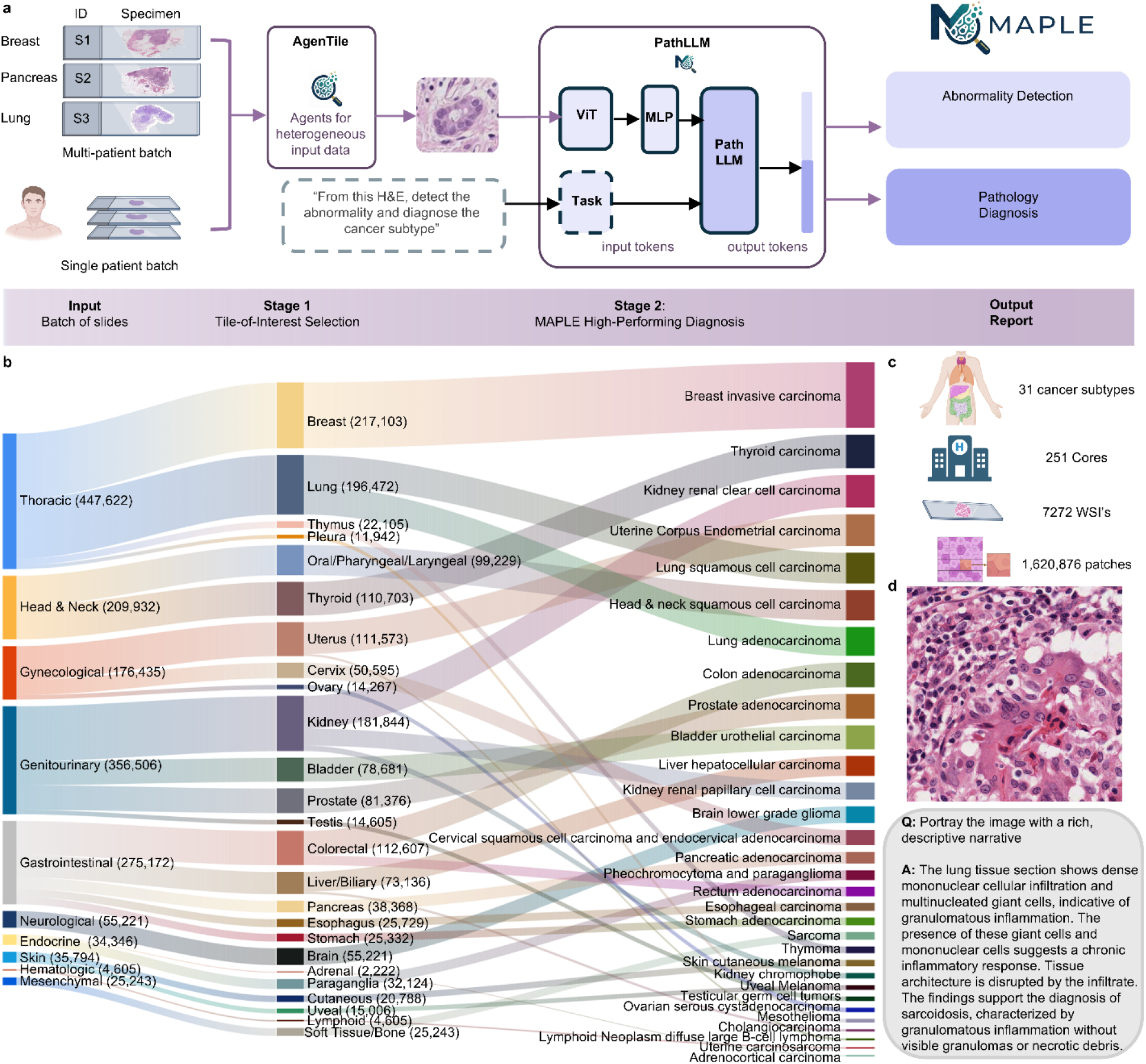
Maples workflow and dataset: **a,** Schematic overview of MAPLE’s hierarchical vision-language architecture. Stage 1: AgenTile performs nuclei instance segmentation and classification at 40x resolution, while the pathology-trained CLIP encoder performs zero-shot tumor classification at downsampled resolution (10x or 5x). A recursive selection function identifies optimal representative patches based on agreement between both modules. Stage 2: PathLLM integrates selected patches (aggregated in groups of 20) with user queries through the pathology-trained CLIP encoder and MLP projection layer, enabling cross-modal reasoning between visual tokens and text inputs within the large language model framework to generate diagnostic reports or targeted responses. **b,** Sankey diagram for oncological cases illustrates the distribution of 1,620,876 training patches across anatomical systems (left), tissue types (center), and 31 cancer subtypes (right). Major categories include thoracic (447,622 patches), gynecological (176,435), genitourinary (356,506), gastrointestinal (275,172), and head and neck (209,932) malignancies. **c,** Dataset composition summary: 31 cancer subtypes, 251 institutional centers, 7,272 whole slide images, and 1,620,876 patches with paired diagnostic text. **d,** Representative H&E-stained histopathology patch depicting lung tissue with dense mononuclear cellular infiltration and multinucleated giant cells characteristic of granulomatous inflammation, supporting a diagnosis of sarcoidosis, illustrating the breadth of pathological conditions captured in the training data beyond oncology.

MAPLE’s workflow accommodates both single-slide analysis and batch processing, supporting either multiple independent patient slides or multiple slides from the same patient queued for overnight processing. The latter mirrors the traditional clinical workflow where resected tissue is split into several representative blocks and histopathology slides are generated from each block and evaluated collectively during diagnosis. MAPLE’s hierarchical architecture enables efficient end-to-end processing from WSI input to structured diagnostic output in 5-10 minutes per slide at 20x magnification and 20-40 minutes at 40x magnification on a single NVIDIA RTX 3090 GPU using automatic patch selection via AgenTile. When patch selection is guided directly by a pathologist, processing time reduces to 1–2 minutes per slide, while maintaining interpretability through selective attention to pathologically relevant regions.

MAPLE’s PathLLM module was trained on a comprehensive dataset comprising 7,272 whole slide images including but not limited to 31 cancer subtypes, sourced from 251 institutional centers to ensure broad representation across clinical settings and patient populations (Fig. 1b and c). Critically, patches were extracted not only from neoplastic regions but also from adjacent non-neoplastic tissue, as well as slides representing a broader spectrum of pathological conditions including infectious diseases, autoimmune conditions, and reactive processes, ensuring the model was exposed to the full histopathological context encountered in clinical practice. The oncological portion of the dataset encompassed major anatomical systems including thoracic (447,622 patches), head and neck (209,932 patches), gynecological (176,435 patches), genitourinary (356,506 patches), gastrointestinal (275,172 patches), neurological (55,221 patches), endocrine (34,346 patches), skin (35,794 patches), hematologic (4,605 patches), and mesenchymal (25,243 patches) malignancies. In total, 1,620,876 patches were extracted and labeled with associated diagnostic text captions, with tissue-level distribution reflecting clinical prevalence: breast (217,103 patches), lung (196,472 patches), kidney (181,844 patches), and colorectal malignancies (112,607 patches), which represent the most abundant categories, alongside substantial representation of prostate malignancies (81,376 patches).

Fig. 1d shows a representative histopathological patch from the training dataset illustrating MAPLE’s exposure to pathological conditions beyond oncology. The image depicts lung tissue with dense mononuclear cellular infiltration and multinucleated giant cells characteristic of granulomatous inflammation, supporting a diagnosis of sarcoidosis. This example highlights that the training dataset encompasses not only neoplastic conditions across 31 cancer subtypes but also non-neoplastic pathological processes including infectious, autoimmune, and inflammatory diseases, equipping MAPLE with the breadth of histopathological knowledge required for comprehensive diagnostic reasoning.

### MAPLE learns discriminative patch-level representations

Accurate diagnostic reasoning in MAPLE depends fundamentally on the quality of patch-level visual representations produced by the image encoder, as morphologically similar tissue types that are not reliably distinguished at this stage cannot be compensated for by downstream language-model reasoning. To evaluate whether MAPLE’s encoder learns diagnostically coherent representations, we isolated it from MAPLE’s pathology-trained CLIP model by detaching the text encoder and benchmarked it against established histopathology vision foundation models, specifically UNI and Virchow, on the BACH breast histopathology dataset (Fig. 2a).^32–34^ Unlike MAPLE’s CLIP-based architecture, UNI and Virchow are standalone vision transformers trained exclusively on histopathology images without contrastive language supervision.^32,33^ The BACH dataset comprises 400 images across four classes (Normal, Benign, In situ carcinoma, and Invasive carcinoma), providing a challenging benchmark for discriminating against benign and malignant breast tissue morphologies.^34^

**Figure 2:**
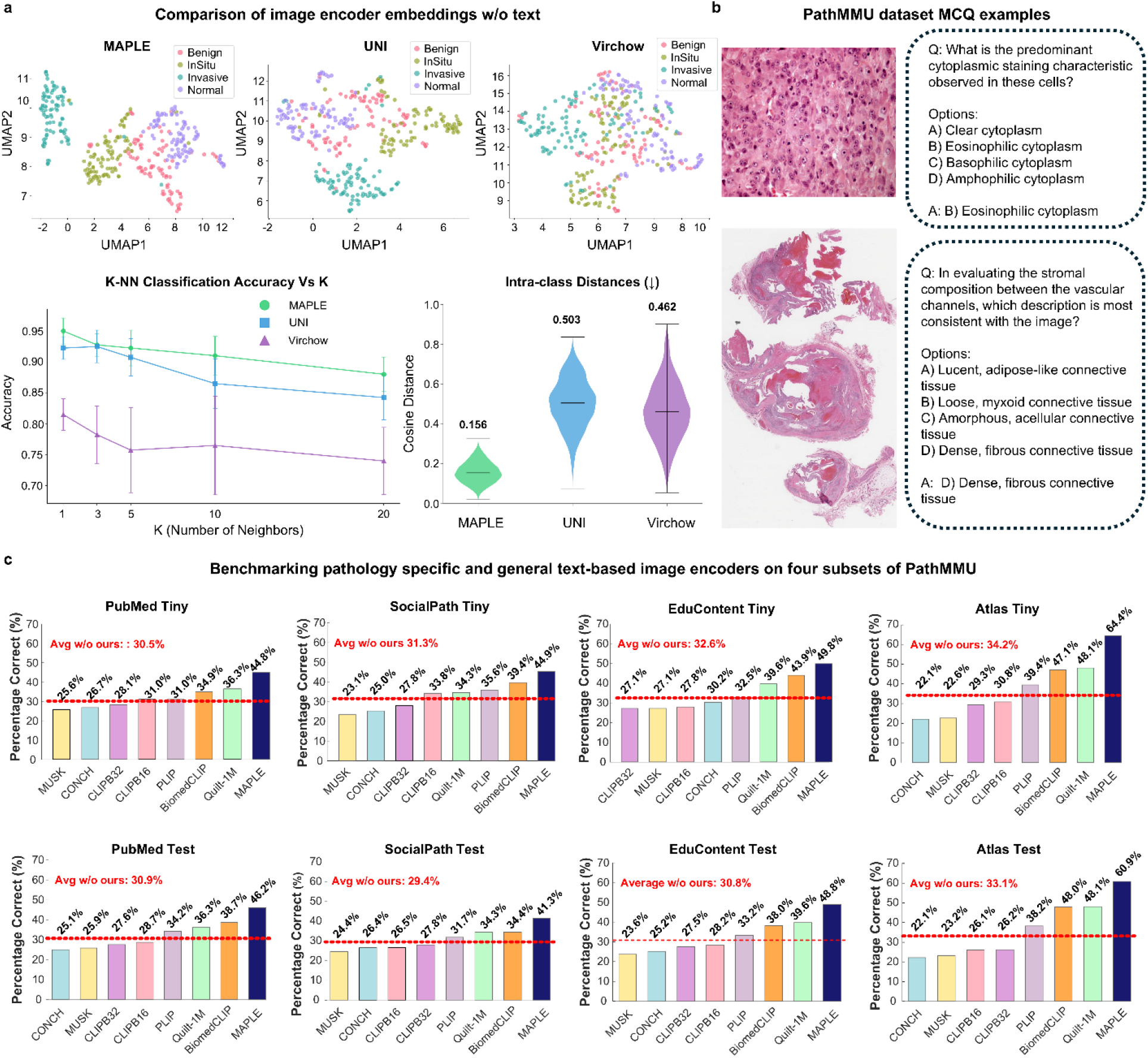
Patch-level evaluation of MAPLE’s image encoder: **a,** Embedding quality comparison of MAPLE’s image encoder (isolated from the CLIP architecture) against UNI and Virchow vision foundation models on the BACH breast histopathology dataset.^32–34^ Top: UMAP projections showing class separation across four diagnostic categories (Normal, Benign, In Situ, Invasive). Bottom left: k-NN classification accuracy as a function of k. Bottom right: Intra-class cosine distance distributions (lower values indicate more compact within-class clustering). **b,** Representative examples from PathMMU evaluation datasets illustrating diverse question types and image sources. Top: H&E-stained histopathology image with a multiple-choice question regarding cytoplasmic staining characteristics. Bottom: Low-magnification tissue architecture assessment of a WSI requiring identification of stromal composition patterns. PathMMU comprises four evaluation domains: PubMed (scientific pathology documents), EduContent (educational YouTube teaching videos), SocialPath (Twitter posts by pathology experts), and Atlas (authoritative pathology textbooks and clinical guidelines).^31^ **c,** Performance of MAPLE’s image encoder across tiny (top) and full (bottom) PathMMU test sets. MAPLE outperformed pathology-specific encoders (CONCH, PLIP, MUSK, Quilt-1M, BiomedCLIP) and general-domain OpenAI CLIP variants across all benchmarks, consistently exceeding baseline averages (red dotted lines).^12–16,37^

UMAP projections of the learned embeddings revealed qualitatively superior class separation for MAPLE compared to UNI and Virchow, with more compact and distinct clusters corresponding to each diagnostic category (Fig. 2a, top).^32,33^ To quantify this observation, we assessed k-nearest neighbor (k-NN) classification accuracy across varying values of k. MAPLE’s image encoder achieved 95.0% accuracy at k=1 (5-fold stratified cross-validation), outperforming both UNI (92.3%) and Virchow (81.5%), with this advantage maintained across all tested k values (Fig. 2a, bottom left). Analysis of embedding space geometry further demonstrated that MAPLE produces more compact within-class representations, with substantially lower intra-class cosine distances (0.156) compared to UNI (0.503) and Virchow (0.462) (Fig. 2a, bottom right).^32,33^ These results indicate that MAPLE’s image encoder learns tightly clustered, diagnostically coherent visual representations that effectively capture morphological differences between pathological categories, outperforming vision-only foundation models on embedding quality metrics. Similar metrics were computed for next-generation models UNI2 and Virchow2, with comparisons to MAPLE presented in Supplementary Fig. 1.^32,35^

To assess generalization of MAPLE’s image encoder to out-of-distribution data, we evaluated its performance on PathMMU, a comprehensive pathologist-validated benchmark comprising multimodal multiple-choice questions spanning diverse pathology domains, including both neoplastic and non-neoplastic conditions. (Fig. 2b).^31^ The PathMMU benchmark includes four subsets used in our evaluation: PubMed (scientific pathology documents), EduContent (educational YouTube teaching videos), SocialPath (Twitter posts by pathology experts), and Atlas (authoritative pathology textbooks and clinical guidelines), totaling 7,691 question-image-answer triplets. A fifth component (PathCLS) was excluded as it is designed for closed-ended visual question answering, whereas MAPLE is optimized for open-ended diagnostic reasoning. For each subset, 200–280 questions ("tiny" split) were independently completed by a panel of seven board-certified pathologists to establish expert-level performance benchmarks. MAPLE’s encoder was trained exclusively on patches obtained from TCGA WSIs and was not exposed to PathMMU images or questions, making this a stringent test of zero-shot generalization and robustness to domain shift across clinical, educational, and informal pathology contexts.^36^

We benchmarked MAPLE’s image encoder against a combination of pathology-specific encoders (CONCH, PILP, MUSK, Quilt-1M, and BiomedCLIP) and general-domain vision encoders (OpenAI CLIP-16, CLIP-32) on PathMMU (Fig. 2c).^12–16,37^ Across both tiny and full test sets, MAPLE consistently outperformed all comparators. On the pathologist-benchmarked tiny subsets, MAPLE achieved 44.8% on PubMed, 44.9% on SocialPath, 49.8% on EduContent, and 64.4% on Atlas, exceeding the individual and average performance of existing pathology-domain encoders on all four datasets (30.5%, 31.3%, 32.6%, and 34.2%, respectively, indicated by red dotted lines) (Fig. 2c, top). Notably, MAPLE surpassed all specialized pathology encoders including MUSK (best: 27.1% on EduContent), CONCH (best: 30.2% on EduContent), PLIP (best: 39.4% on Atlas), BiomedCLIP (best: 47.1% on Atlas), and Quilt-1M (best: 48.1% on Atlas), as well as the original OpenAI CLIP variants (CLIP-16 and CLIP-32, ranging from 27.1% to 33.8%) (Fig. 2c, top).

Performance gains were maintained on the full test sets (Fig. 2c, bottom), where MAPLE achieved 46.2% on PubMed, 41.3% on SocialPath, 48.8% on EduContent, and 60.9% on Atlas, consistently exceeding baseline averages of existing pathology-domain encoders (30.9%, 29.4%, 30.8%, and 33.1%, respectively, indicated by red dotted lines). Margins over pathology-specific encoders ranged from 16.9 to 38.8 percentage points (Fig. 2c, bottom). These results establish MAPLE’s image encoder as a new state-of-the-art for pathology image representation learning at the patch level.

### Zero-shot classification performance of MAPLE

We next evaluated the zero-shot classification capabilities of MAPLE’s image encoder across diverse histopathology tasks, as the ability to classify tissue types without task-specific fine-tuning is essential for practical deployment across the broad range of diagnostic entities encountered in clinical practice, where annotated training data for every possible condition cannot be assumed. MAPLE demonstrated state-of-the-art performance across two out of three public benchmarks, outperforming pathology-specific vision encoders (Fig. 3a). On BACH, a four-class breast histopathology dataset distinguishing normal, benign, in situ carcinoma, and invasive carcinoma, MAPLE achieved 78.3% balanced accuracy and 78.1% macro F1 score, exceeding the next-best performers (CONCH: 64.7% balanced accuracy, and MUSK: 61.5% macro F1) by 13.6 to 16.6 percentage points (Fig. 3a).^34^ On PatchCamelyon, a binary lymph node metastasis detection dataset, MAPLE attained 85.0% balanced accuracy and 84.9% macro F1, surpassing CONCH (71.0% and 70.6%), PLIP (71.0% and 71.0%), and outperforming BiomedCLIP (75.5% and 75.5%) and Quilt-1M (77.0% and 77.0%), while performing comparably to MUSK (85.3% and 85.3%) (Fig. 3a).^38^

**Figure 3:**
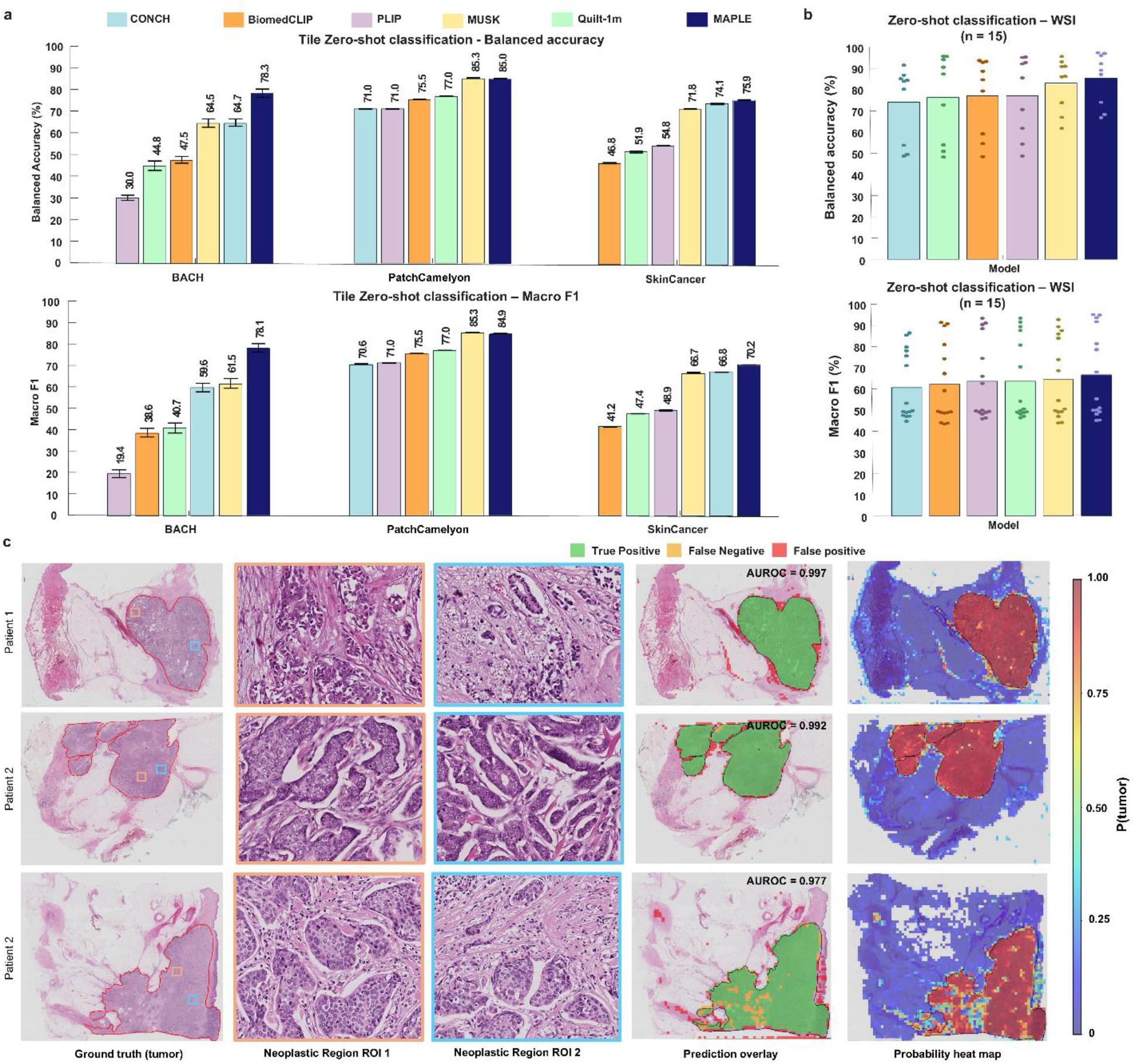
Zero-shot classification performance. **a,** Zero-shot classification performance of MAPLE’s image encoder compared to pathology-specific encoders (CONCH, PLIP, MUSK, Quilt-1M, BiomedCLIP) across three public histopathology datasets.^12–16^ Top: Balanced accuracy across BACH (four-class breast histopathology), PatchCamelyon (binary lymph node metastasis detection), and SkinCancer (16-class dermatological conditions).^34,38,39^ Bottom: Corresponding macro F1 scores. **b,** Zero-shot classification performance on clinical breast tissue whole-slide images (n = 15), validated against annotations from a board-certified pathologist at Johns Hopkins. Top: Balanced accuracy. Bottom: Macro F1 score. **c,** Spatial visualization of patch-level predictions across three representative cases. From left to right: H&E-stained tissue sections with ground truth tumor annotations, magnified neoplastic regions of interest (ROI 1 and ROI 2), prediction overlay showing tumor (green), false negatives (yellow), and false positives (red), and probability heatmaps showing continuous tumor likelihood scores (red = high probability, blue = low probability). MAPLE achieved AUROC values of 0.997, 0.992, and 0.977 for the three cases shown.

For the multi-class SkinCancer dataset encompassing 16 dermatological conditions, MAPLE reached 75.9% balanced accuracy and 70.2% macro F1, outperforming BiomedCLIP (46.8% and 41.2%), Quilt-1M (51.9% and 47.4%), PLIP (54.8% and 48.9%), MUSK (71.8% and 66.7%), and CONCH (74.1% and 66.8%) (Fig. 3a).^39^

In the pathologist-annotated clinical WSI breast sample cohort (Fig. 3b), MAPLE achieved superior performance with 85.3% balanced accuracy and 66.5% macro F1 for tumor detection (Fig. 3b), outperforming CONCH (73.7% and 60.8%), Quilt-1M (76.5% and 63.7%), BiomedCLIP (77.1% and 63.6%), PLIP (77.3% and 63.7%), and MUSK (82.8% and 64.7%) (Fig. 3b). Spatial visualization of patch-level predictions demonstrated MAPLE’s capacity for accurate tumor localization across diverse histological presentations, with predicted boundaries closely matching pathologist annotations (Fig. 3c).

These results establish MAPLE’s vision encoder as capable of zero-shot generalization across histopathology domains, achieving expert-level discrimination without task-specific training. This capability is critical for deployment across the diverse diagnostic scenarios encountered in clinical pathology practice.

### MAPLE’s multimodal reasoning exceeds pathologist performance

Having established that MAPLE’s image encoder learns superior patch-level representations (Figs. 2-3), we next evaluated the complete PathLLM pipeline, integrating the image encoder with the large language model, on multimodal reasoning tasks. We benchmarked MAPLE against state-of-the-art multimodal models and board-certified pathologists on PathMMU (Fig. 4a).

**Figure 4.**
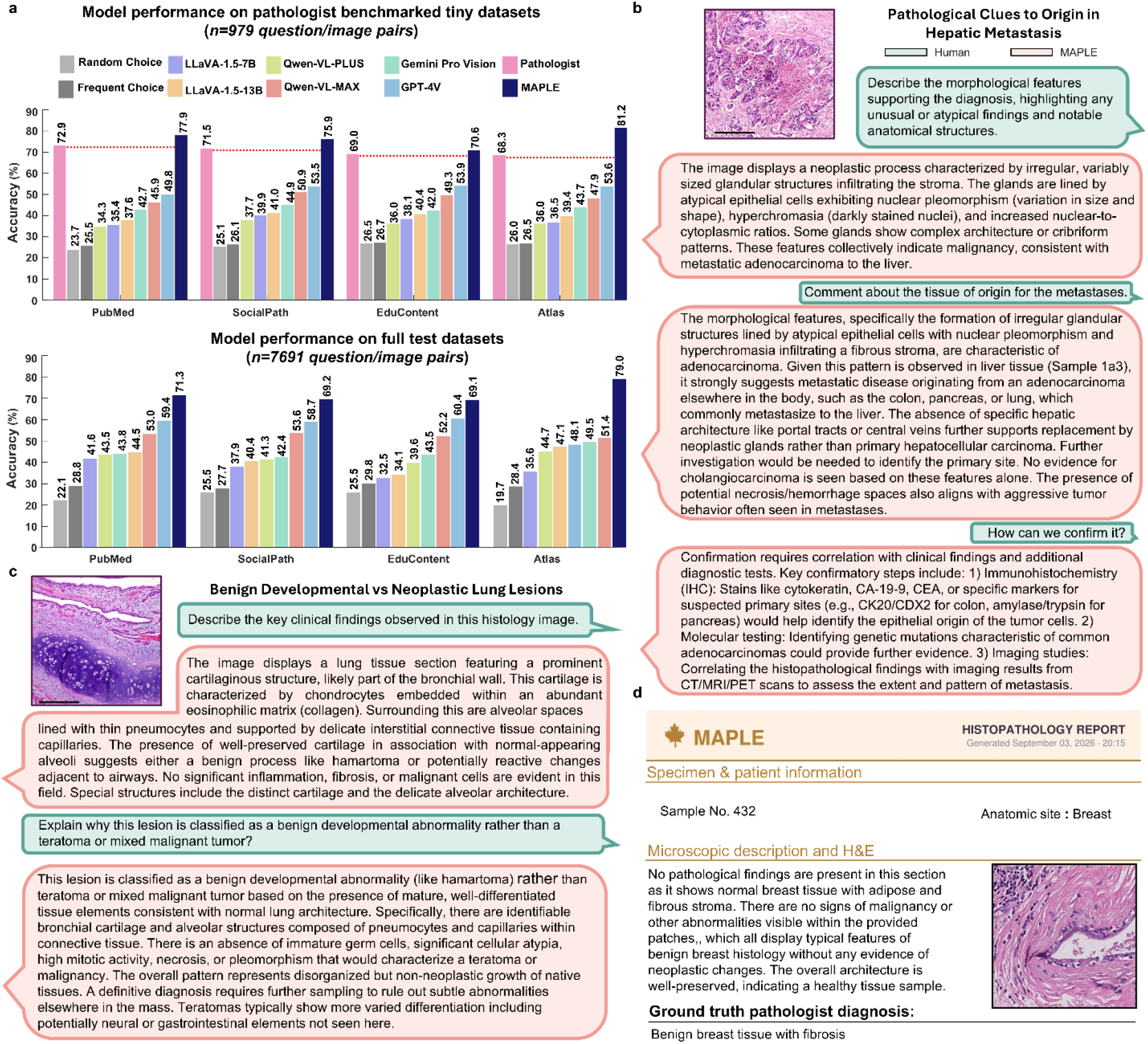
MAPLE’s multimodal diagnostic performance, clinical reasoning, and report generation. **a,** Performance of MAPLE on the four PathMMU domains (PubMed, SocialPath, EduContent, and Atlas), compared with state-of-the-art vision-language models and expert pathologists. Top: Multiple choice question (MCQ) accuracy on the pathologist-benchmarked tiny subjects (n = 979 image-question pairs), with pink bars and dashed reference lines indicating average pathologist performance. MAPLE outperformed the evaluated models and the pathologist benchmark across all four domains. Bottom: MCQ accuracy on the full test sets (n = 7,691 image-question pairs), demonstrating MAPLE’s performance relative to competing multimodal models. **b,** Representative multi-turn diagnostic dialogues illustrating MAPLE’s ability to interpret histopathological findings and provide context-dependent clinical reasoning. In a case of hepatic metastasis, MAPLE identifies morphological features consistent with metastatic adenocarcinoma, discusses potential primary sites, and recommends ancillary investigations to resolve diagnostic uncertainty. Human prompts are shown in green and MAPLE responses in red. **c,** In a case of pulmonary hamartoma, the model encounters cartilaginous tissue within lung parenchyma, a finding that may superficially mimic metastasis or teratoma. Rather than reflexively attributing malignancy, it evaluates the absence of cytological atypia, architectural organization, and lack of infiltrative growth to correctly classify the lesion as benign, demonstrating that morphological heterogeneity alone does not inherently indicate malignancy. **d,** Representative MAPLE-generated histopathology report for a benign breast tissue specimen, illustrating the integration of specimen information, microscopy description, and histological imaging. The generated diagnosis is consistent with the pathologist’s reference diagnosis of benign breast tissue with fibrosis.

On the pathologist-benchmarked subsets (n = 979 question-image pairs), MAPLE achieved superior performance across all four evaluation domains, representing the first multimodal model to surpass pathologist performance across PathMMU benchmarks and substantially exceeding state-of-the-art multimodal models (Fig. 4a, top). On PubMed, MAPLE attained 77.9% accuracy, outperforming the pathologist average (72.9%) by 5 percentage points and far exceeding GPT-4V (49.8%) by 28.1 points (Fig. 4a, top). Similarly, models of comparable size, including LLaVA-1.5-13B (37.6%), performed substantially lower. On SocialPath, MAPLE achieved 75.9% accuracy compared to 71.5% for pathologists, with notable margins over GPT-4V (53.5%), Qwen-VL-MAX (50.9%), and LLaVA-1.5-13B (41.0%) (Fig. 4a, top). On EduContent, MAPLE scored 70.6%, exceeding the pathologist average of 69.0% and outperforming GPT-4V (53.9%), Qwen-VL-MAX (49.3%), Gemini Pro Vision (42.0%), and LLaVA-1.5-13B (40.4%) (Fig. 4a, top).

MAPLE’s advantage was most pronounced on Atlas, a dataset comprising questions from authoritative pathology textbooks and clinical guidelines. MAPLE reached 81.2% accuracy, 12.9 points above the pathologist benchmark (68.3%), while GPT-4V achieved only 53.6% and other larger models including Qwen-VL-MAX (47.9%) and Qwen-VL-PLUS (36.0%) remained below human expert performance (Fig. 4a, top).

Performance trends on the full test sets (n = 7,691 question-image pairs) mirrored those observed in the pathologist-benchmarked subsets, with MAPLE maintaining substantial advantages across all domains (Fig. 4a, bottom). MAPLE achieved 71.3% on PubMed, 69.2% on SocialPath, 69.1% on EduContent, and 79.0% on Atlas (Fig. 4a, bottom), consistently outperforming GPT-4V by margins of 8.7 to 30.9 percentage points. Compared to similarly sized models, MAPLE exceeded LLaVA-1.5-13B by 26.8 to 35 percentage points across benchmarks and surpassed larger architectures including Qwen-VL-MAX (best: 53.6% on SocialPath) and Gemini Pro Vision (best: 49.5% on Atlas) (Fig. 4a, bottom).

Beyond quantitative performance, MAPLE demonstrates sophisticated clinically coherent diagnostic reasoning consistent with conventional pathologic assessment (Fig. 4b and c). In a representative case of hepatic metastasis, when asked to describe morphological features, MAPLE correctly identified irregular, variably sized glandular structures infiltrating the stroma, lined by atypical epithelial cells exhibiting nuclear pleomorphism, hyperchromasia, and increased nuclear-to-cytoplasmic ratios, features collectively indicating metastatic adenocarcinoma (Fig. 4b). When prompted about tissue of origin, MAPLE generated a ranked differential (colon, pancreas, or lung), synthesizing morphological cues with epidemiological patterns of hepatic metastasis. It correctly ruled out primary hepatocellular carcinoma by noting absent native hepatic architecture (portal tracts, central veins), indicating neoplastic replacement (Fig. 4b). MAPLE acknowledged that distinguishing among adenocarcinoma primaries requires immunohistochemical confirmation, demonstrating calibrated confidence where morphology provides probabilistic guidance rather than definitive answers. When asked to confirm the probable tissue of origin, MAPLE recommended ancillary steps: it appropriately recommended CK20/CDX2 for colon immunohistochemistry, but inappropriately recommended amylase/trypsin immunohistochemistry for pancreas, and appropriately recommended molecular testing for characteristic mutations, and imaging correlation (Fig. 4b).

This calibrated uncertainty reiterated with the pulmonary hamartoma case, where MAPLE demonstrated confident classification (Fig. 4c). Upon encountering cartilaginous tissue within lung parenchyma, a finding that could superficially trigger metastatic suspicion, MAPLE correctly identified chondrocytes embedded within eosinophilic matrix surrounded by well-preserved alveolar architecture (Fig. 4c). When challenged to explain why this represents benign developmental abnormality rather than teratoma or mixed malignant tumor, MAPLE reasoned systematically: mature, well-differentiated tissue elements consistent with normal lung architecture; absence of immature germ cells, significant cellular atypia, high mitotic activity, necrosis, or pleomorphism that would characterize a teratoma or malignancy; and lack of varied differentiation (neural, gastrointestinal elements) expected in teratoma (Fig. 4c). This contextual tissue discrimination, recognizing that "different tissue" does not equal "dangerous tissue," reflects sophisticated reasoning beyond pattern-matching. Unlike metastatic deposits, which are characterized by nuclear pleomorphism, mitotic activity, and infiltrative destruction of native tissue architecture, hamartomatous lesions comprise mature tissue elements without nuclear atypia or infiltrative growth, a diagnostically demanding distinction requiring recognition of cellular composition, nuclear morphology, and tissue of origin that MAPLE resolves correctly through integrated morphological reasoning. This ability to confidently classify benign lesions was further demonstrated in a breast tissue case (Fig. 4d), where MAPLE correctly recognized normal lobular architecture and stromal fibrosis without atypical features, and organized these findings into a structured histopathology report whose final diagnosis matched the ground truth diagnosis established by a board-certified breast pathology specialist at Johns Hopkins Hospital

This calibrated reasoning, distinguishing benign from malignant findings while exercising appropriate uncertainty, was further validated by the perineural invasion case (Supplementary Fig. 2b). Here, MAPLE correctly recognized tumor cells tracking along nerve sheaths as clinically significant invasion rather than incidental anatomical proximity, demonstrating its ability to avoid both over-interpretation of benign findings and under-recognition of true pathology. Additional supplementary cases illustrate complementary reasoning domains: clinicopathological integration correlating leiomyoma histology with menstrual symptomatology when provided with patient history (Supplementary Fig. 2a); diagnostic robustness through guardrail testing (Supplementary Fig. 2c); HER2 scoring despite limited immunohistochemistry representation in training data (Supplementary Fig. 3a and b); grading competency through Gleason and ISUP system application in prostatic adenocarcinoma (Supplementary Fig. 3c and d) and clear cell renal cell carcinoma (Supplementary Fig. 4c); and therapeutic reasoning in uterine sarcoma management (Supplementary Fig. 4a).

Beyond oncological diagnosis, MAPLE demonstrated diagnostic breadth across infectious and autoimmune disease categories, correctly identifying Strongyloides infection (Supplementary Fig. 5a), cryptococcal meningitis (Supplementary Fig. 5b), Hashimoto’s thyroiditis (Supplementary Fig. 5c and d), and autoimmune hepatitis (Supplementary Fig. 5e), illustrating the model’s capacity to generalize across histopathological disease domains without task-specific fine-tuning.

Collectively, these cases demonstrate MAPLE’s capacity for hierarchical reasoning, progressing from morphological description through differential diagnosis to actionable recommendations.

### AgenTile performs cellular-level triage to enable whole-slide analysis

Clinical pathology diagnosis requires cellular-level assessment, identifying neoplastic cells, characterizing nuclear morphology, and evaluating tissue architecture. However, gigapixel WSIs contain millions of cells, making exhaustive analysis computationally prohibitive, and indiscriminate patch sampling risks presenting the language model with uninformative or artefactual regions, diluting diagnostic signal and reducing reasoning quality. By selectively triaging only the most diagnostically relevant regions prior to multimodal reasoning, MAPLE ensures that PathLLM’s limited context window is allocated to patches most likely to contain actionable pathological findings. MAPLE addresses this through AgenTile, a lightweight vision transformer that performs rapid nuclei segmentation and classification at 40x resolution, enabling intelligent triage of diagnostically relevant regions before resource-intensive multimodal reasoning (Fig. 5a).

**Figure 5:**
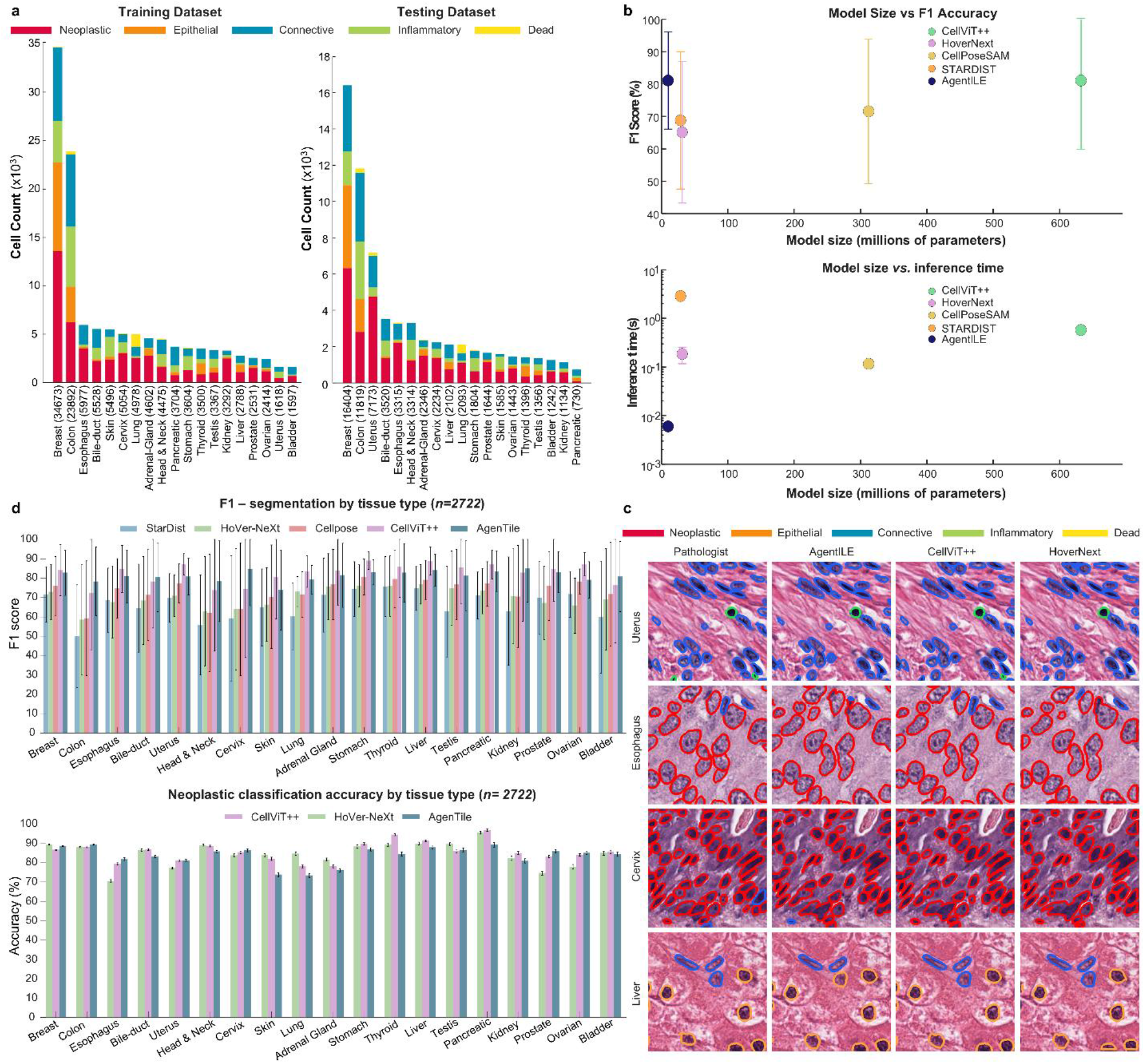
AgenTile for high resolution gigapixel WSI processing via cell segmentation and classification for neoplastic patch selection. **a,** Distribution of annotated nuclei across 19 tissue types in the PanNuke dataset, stratified by cell category (neoplastic, epithelial, connective, inflammatory, dead) for training (left) and testing (right) sets. Tissue representation varied from high (breast, colon, esophagus) to limited (bladder, ovarian, pancreatic) across both splits, with all five cell categories present within each tissue type. **b,** Comparative analysis of model size versus performance (top) and inference time (bottom) across nuclei segmentation architectures. AgenTile (∼10 million parameters) achieved F1 segmentation accuracy of 81.1%, matching CellViT++ (632 million parameters) while providing a ∼65-fold reduction in model size and 95-fold speedup in inference time (0.006s vs 0.58s per patch on an NVIDIA RTX 3090 GPU). Other benchmarked models: CellposeSAM (0.16s, F1=71.6), StarDist (0.19s, F1=65.1), and HoverNext (2.94s, F1=68.8). **c,** Qualitative comparison of nuclei segmentation and classification across representative tissue types (uterus, esophagus, cervix, liver) comparing pathologist annotations, AgenTile, CellViT++, and HoverNext. Cell categories are color-coded: neoplastic (red), epithelial (orange), connective (blue), inflammatory (green), dead (yellow). **d,** Per-tissue F1 segmentation scores and neoplastic classification accuracy across 19 tissue types for all benchmarked models. AgenTile achieved the highest segmentation F1 in 6 of 19 tissues compared to CellViT++ and outperformed StarDist, HoverNext, and CellposeSAM across all 19 tissues, while exceeding CellViT++ in neoplastic classification accuracy in 8 of 19 tissue types.

AgenTile operates as the first stage of MAPLE’s hierarchical pipeline, scanning WSIs to identify patches enriched for neoplastic cells, inflammatory infiltrates, or other features warranting detailed analysis. By quantifying cellular composition across the slide, AgenTile provides spatial context for MAPLE’s recursive patch selection algorithm, which integrates 40x cellular features from AgenTile with 10x and 5x tissue architectural patterns extracted by the CLIP image encoder to identify the most diagnostically informative regions.

AgenTile was trained on the pathologist-validated PanNuke dataset, which comprises 7,901 image patches (256x256 pixels, 40x magnification, 0.25 µm/pixel) with 189,744 annotated nuclei across 19 tissue types and five cell categories (neoplastic, epithelial, connective, inflammatory, and dead cells).^40^ Following the predefined three-fold partitioning of PanNuke, two folds were used for training and one for testing, with tissue representation varying from high (breast, colon, esophagus) to limited (bladder, ovarian, pancreatic) across both splits (Fig. 5a).

We benchmarked AgenTile against established segmentation models spanning different architectural paradigms: ViT-based (CellViT++, CellposeSAM), CNN-based (HoverNext), and U-Net-based (StarDist). AgenTile achieved a global F1 segmentation score of 81.1 (Fig. 5b), significantly outperforming CellposeSAM, StarDist, and HoverNext (71.6, 65.1, and 68.8, respectively) and matching CellViT++ performance, while requiring only ∼10 million parameters versus 632 million for CellViT++ (∼65-fold fewer parameter) (Fig. 5b).^41–44^ This efficiency translated to substantially faster inference: 0.006s per patch for AgenTile versus 0.58s for CellViT++ (∼95-fold speedup), with other models requiring 0.12s (CellposeSAM), 0.19s (StarDist), and 2.88s (HoverNext) per patch on an NVIDIA 3090 GPU (Fig. 5b).

Qualitative comparison across representative tissue types (uterus, esophagus, cervix, liver) demonstrated comparable segmentation fidelity across AgenTile, CellViT++, and HoverNext, with all methods closely approximating pathologist annotations (Fig. 5c). In per-tissue quantitative analysis, AgenTile achieved the highest segmentation F1 scores in 6 of 19 tissues compared to CellViT++ and outperformed StarDist, HoverNext, and CellposeSAM across all 19 tissues. For neoplastic classification of segmented cells, AgenTile exceeded CellViT++ in 8 of 19 tissue types and HoverNext in 6 (Fig. 5d), while providing the fastest inference time among all benchmarked approaches (Supplementary Fig. 6a).

### MAPLE’s streamlined Graphical User Interface (GUI) for clinical operational needs

To facilitate clinical translation, we developed a unified graphical user interface integrating MAPLE’s complete hierarchical workflow, from whole-slide image processing through diagnostic report generation (Fig. 6). The generated report provides a succinct microscopic description summarizing key histopathological findings, accompanied by spatial visualization of suspicious regions identified as neoplastic and a probabilistic tumor mapping that displays attention-weighted likelihood of malignancy across the slide (Fig. 6a). This multimodal output format mirrors conventional pathology reporting standards while augmenting them with comprehensive spatial visualization, supporting both clinical decision-making and quality assurance review.

**Figure 6.**
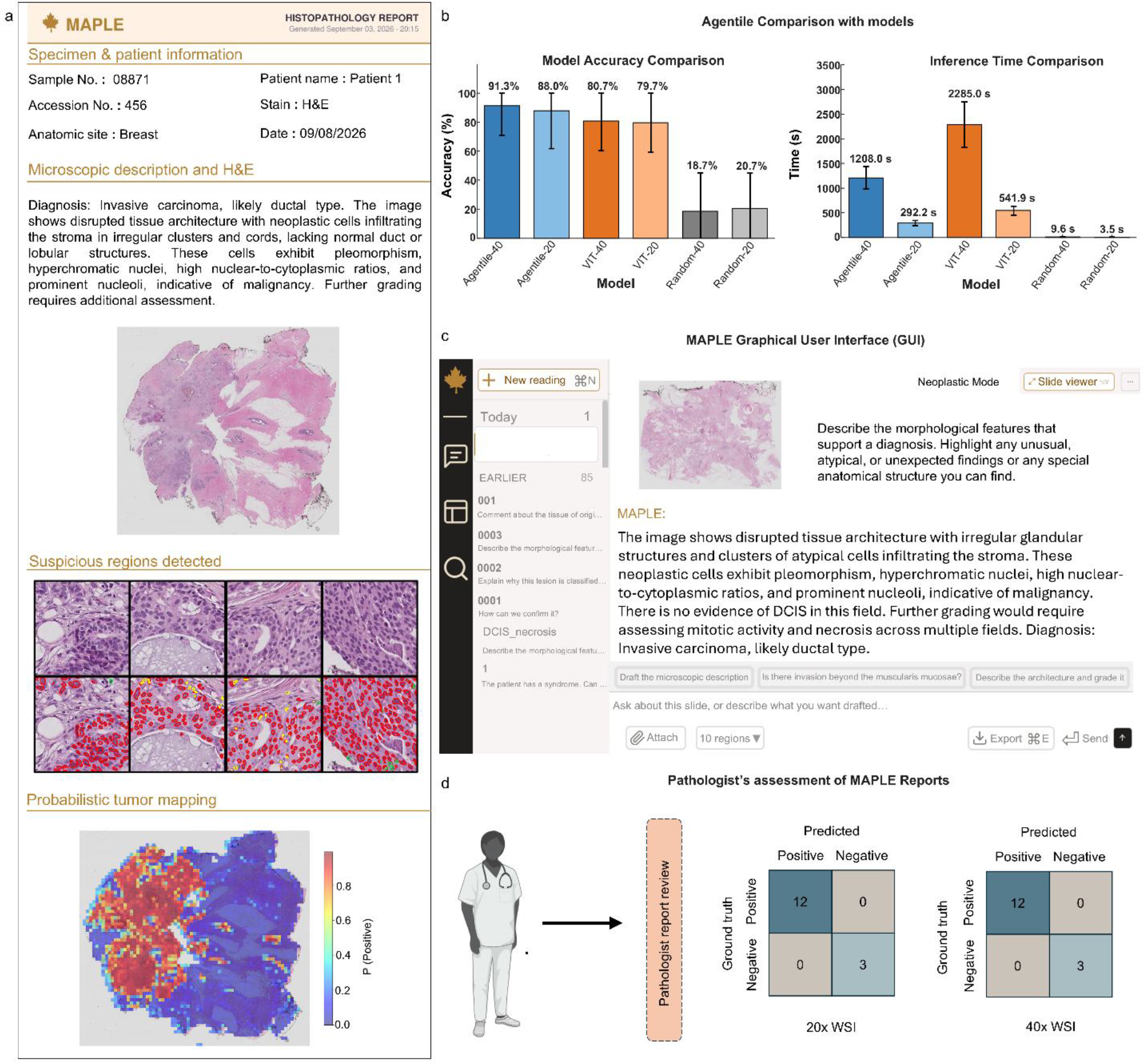
Clinical deployment, ablation analysis, and whole-slide diagnostic validation. **a,** Comprehensive diagnostic report generation module producing structured outputs with case metadata (patient identifiers, specimen details, clinical history), narrative microscopic description with histopathological findings, spatially-annotated suspicious regions demarcated as neoplastic, and probabilistic tumor mapping displaying attention-weighted likelihood of malignancy across the slide. **b,** Ablation analysis comparing AgenTile-guided patch selection versus standard ViT-based selection. Model accuracy comparison (left) demonstrates AgenTile’s superior performance against the ViT at both 40x (91.1% vs 80.7%) and 20x (88.0% vs 79.7%) magnifications, attributable to cellular-level neoplastic classification capabilities. Inference time comparison (right) shows AgenTile’s computational efficiency, completing patch selection in approximately 1208s at 40x versus 2285s for standard ViT (1.9-fold speedup). **c,** Graphical user interface demonstrating batch whole-slide image processing and conversational capabilities. The neoplastic mode toggle enables selection between AgenTile-guided intelligent sampling (prioritizing diagnostically relevant regions) or user-guided sampling. Users can interactively query the system with natural language questions regarding histopathological features, differential diagnoses, or morphological interpretations. **d,** Whole-slide diagnostic validation on 15-WSI breast tissue cohort against ground truth diagnoses established by board-certified breast pathology specialist. Confusion matrices show performance at 20x (left) and 40x (right) magnifications, with 100% accuracy (15/15 correct) at both resolutions and no false positives at 40x.

Additionally, we evaluated the contribution of AgenTile’s cellular-level triage to overall diagnostic accuracy through ablation analysis (Fig. 6b). Comparing AgenTile-guided patch selection against standard ViT-based selection at matched magnifications, AgenTile achieved higher accuracy at both 40x (91.3% vs 80.7%) and 20x (88.0% vs 79.7%), demonstrating that cellular classification capabilities for detecting neoplastic regions critically inform downstream diagnostic performance. AgenTile also provided dramatically faster inference, completing patch selection in approximately 1200s at 40x compared to 2285s for the ViT (1.9-fold speedup), with similar gains at 20x (Fig. 6b, right).

The system enables batch processing of multiple WSIs or pre-extracted patches for high-throughput diagnostic screening, representing the first pathology-domain model to accommodate this capability (Fig. 6c). Batch processing mirrors routine clinical workflows where multiple slides (typically 5-10 per case) are evaluated simultaneously for comprehensive diagnosis. A configurable neoplastic mode toggle allows users to select between AgenTile-guided patch selection, prioritizing regions with abnormal cellular architecture, or user-guided region selection for scenarios where regions of interest have been pre-identified by pathologists.

Beyond diagnostic workflows, MAPLE functions as a standalone pathology-focused conversational chatbot for educational and training applications (Fig. 6c). Users can iteratively query the system with natural language questions regarding histopathological features, differential diagnoses, or morphological interpretations, with output that provides spatial visualization and explainable reasoning traces.

We validated MAPLE’s whole-slide diagnostic accuracy on the 15-WSI breast tissue cohort against ground truth diagnoses established by a board-certified breast pathology specialist at Johns Hopkins Hospital (Fig. 6d). The cohort comprised 9 invasive carcinoma cases, 3 cases of low-grade ductal carcinoma in situ, and 3 benign cases, representing a wide spectrum of breast cancer pathology encountered in routine clinical practice. At both 40x and 20x magnification, MAPLE correctly diagnosed 15 of 15 cases (100.0% accuracy), demonstrating consistent performance across magnifications. These results establish MAPLE’s readiness for clinical piloting as a diagnostic support tool, with performance approaching specialist-level accuracy while providing scalable and reproducible interpretations.

## DISCUSSION

Computational pathology has advanced substantially through deep learning models trained on histopathological images,^45–50^ yet clinical translation has been hindered by three fundamental limitations. First, existing systems are unable to process gigapixel-scale whole slide images at full diagnostic resolution, the operational standard in digital pathology, within the memory and computational constraints of a single GPU, which represents the hardware most commonly available in clinical settings, thereby severely limiting practical deployability.^27,51–54^ Second, while several models incorporate natural language interfaces, their outputs are oriented toward research and engineering terminology rather than the structured, diagnostically precise language familiar to practicing pathologists, reducing their utility at the clinical interface.^55–57^ Third, comprehensive benchmarking of model performance against expert pathologist standards has been lacking; most systems are evaluated on narrow task-specific metrics that neither reflect the breadth of real-world diagnostic reasoning nor establish whether models approach, let alone surpass, expert-level performance.^58–60^ Taken together, these gaps have prevented the emergence of a computationally and clinically fluent pathology AI systems suitable for routine deployment as a copilot in existing clinical workflows.

Here we introduce MAPLE, a Multimodal Assistant for Pathology vision-Language Evaluation, whose hierarchical architecture addresses these limitations by combining selective patch-level tissue abnormality detection with multimodal diagnostic reasoning. AgenTile processes whole slide images at full 40x resolution to identify diagnostically relevant regions at the cellular level, aided by a coarse tumor probability heatmap generated by MAPLE’s image encoder from downsampled whole slide images. These two signals are combined to guide patch selection, directing focus toward regions of highest diagnostic relevance before the selected patches are passed to PathLLM for vision-language integration and report generation. This two-stage design enables comprehensive WSI assessment while remaining computationally tractable on a single consumer-grade GPU. AgenTile is approximately 65-fold smaller and approximately 95-fold faster than the leading cell-segmentation benchmark, CellViT++, restoring the capacity for holistic slide-level diagnosis comparable to pathologist review as performed in current clinical workflows, without requiring specialized infrastructure.

MAPLE’s pathology-adapted image encoder demonstrated strong zero-shot classification performance across three public benchmarks and clinical WSI cohorts, outperforming both domain-specific models and general-purpose encoders. As a complete multimodal system, PathLLM, trained on 7,272 whole slide images spanning 31 cancer subtypes from 251 institutional centers, achieved competitive accuracy on PathMMU’s full test sets spanning scientific pathology literature (PubMed), educational video content (EduContent), expert pathologist social media posts (SocialPath), and authoritative pathology textbooks and clinical guidelines (Atlas), demonstrating robust generalization across acquisition protocols, staining variations, and tissue types that narrower task-specific classifiers have struggled to match. On PathMMU benchmark subsets where pathologist consensus scores are available, MAPLE is the first model to exceed pathologist-level performance across all four evaluated domains, a result that reflects the breadth and diversity of its training data and, while confined to these structured subsets rather than open-ended clinical tasks, nonetheless illustrates the model’s capacity for expert-level domain knowledge in question-answering contexts. Notably, these results were achieved with a 14-billion parameter architecture, comparing favorably against substantially larger industry-built systems with estimated parameter counts orders of magnitude greater, suggesting that domain-adapted training is a more important determinant of diagnostic performance than raw model scale, and that a system achieving competitive accuracy at a fraction of the computational cost is more likely to be integrated into routine institutional workflows than one requiring specialized infrastructure.

MAPLE generates reports integrating structured findings, narrative impressions with morphological reasoning, annotated representative regions, and whole-slide probability heatmaps. This output format is designed to align with conventional pathology reporting while providing spatial-visual-textual integration to support pathologist review. The illustrated clinical reasoning examples demonstrate qualitative interpretive capability; systematic evaluation of reasoning consistency and uncertainty behavior across larger case cohorts.

MAPLE is designed as a copilot to augment, not replace, pathologist expertise. The system’s intended deployment model involves batch processing of incoming cases, generating preliminary reports for pathologist to integrate into their evaluations. The integrated interface supports simultaneous processing of multiple whole slide images, mirroring routine clinical workflows. This approach addresses critical bottlenecks in high-throughput settings, screening large caseloads, triaging complex cases, and reducing turnaround times, while preserving pathologist decisional authority. Pathologists retain full oversight, using MAPLE’s outputs as a structured second opinion that highlights regions of concern, suggests differential diagnoses, and provides morphological reasoning for consideration.

In a preliminary clinical validation on 15 breast whole slide images annotated by board-certified specialists, MAPLE achieved 100% diagnostic accuracy (15/15), correctly classifying both malignant and benign cases across the cohort. These results are encouraging but should be interpreted cautiously: the cohort is small, drawn from a single institution, and not representative of the full range of breast pathology encountered in practice. Independent, multi-institutional prospective validation will be necessary to establish generalization to diverse acquisition platforms, patient populations, and subspecialty contexts.

Several limitations must be acknowledged. First, MAPLE requires high-quality digitized WSIs at 40x or 20x magnification; lower-resolution scans or significant tissue artifacts can propagate through the pipeline and affect diagnostic accuracy. Second, while AgenTile was trained across 19 tissue types, performance may vary for rare tumor subtypes underrepresented in the training data. Third, diagnostic features occupying minimal slide area may be missed if not captured during patch selection, although configurable sampling modes, including expert-guided patch selection, partially mitigate this risk. Fourth, MAPLE cannot incorporate clinical context beyond the provided query: patient history, prior diagnoses, and ancillary testing results are routinely integrated by pathologists and remain outside the current system’s scope, underscoring the necessity of human oversight. Finally, and most importantly, independent multi-institutional clinical validation across diverse subspecialties, scanner platforms, and staining protocols will be essential before deployment in real-world diagnostic workflows. Systematic evaluation of diagnostic reasoning quality, beyond the qualitative examples presented here, represents a further necessary step toward responsible clinical translation.

In conclusion, MAPLE advances computational pathology by integrating hierarchical multiscale tissue analysis with natural language reasoning in an architecture designed for practical institutional deployment. Rather than requiring extreme computational resources or accepting substantially lower accuracy, MAPLE achieves expert-competitive performance within a parameter-efficient framework. Future work will expand the system to incorporate immunohistochemical stains, molecular annotations, and electronic health record integration for fully contextualized diagnostic support, alongside prospective multi-institutional validation to establish generalization across diverse clinical environments. MAPLE represents a step toward scalable, interpretable AI tools that enhance pathologist productivity and diagnostic consistency while maintaining the primacy of human expertise in patient care.

## FIGURES AND CAPTIONS

**Supplemental Fig 1.**
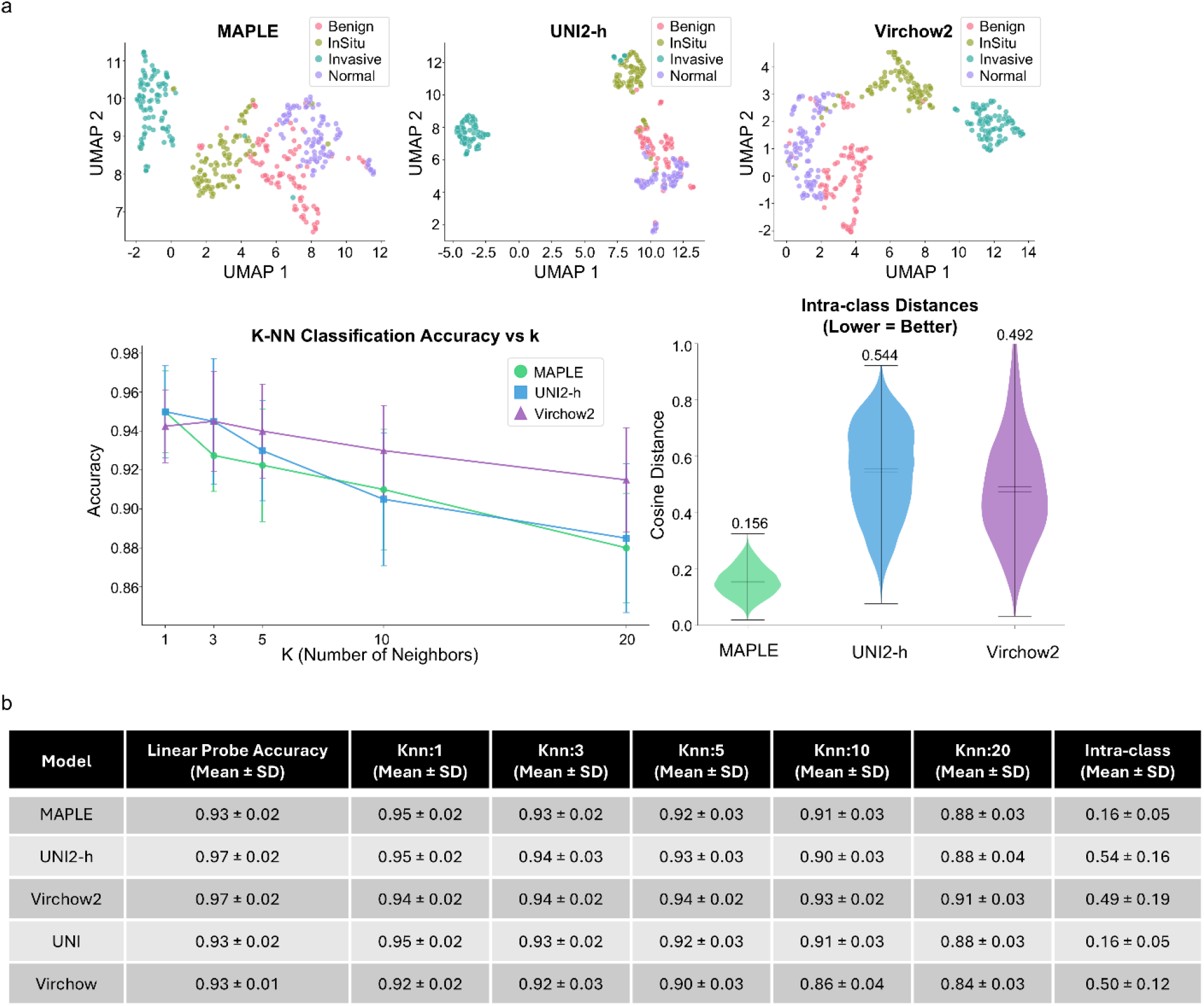
Comparison of patch-level image representations on the BACH dataset. **a,** UMAP projections of MAPLE, UNI2-h, and Virchow2 embeddings across four diagnostic categories from the BACH dataset, together with k-nearest neighbor (k-NN) classification accuracy and intra-class cosine distance distributions. **b,** Comparison of linear-probe accuracy, k-NN classification accuracy, and intra-class cosine distances across five image encoders. Values are reported as mean ± standard deviation (SD).

**Supplemental Fig 2.**
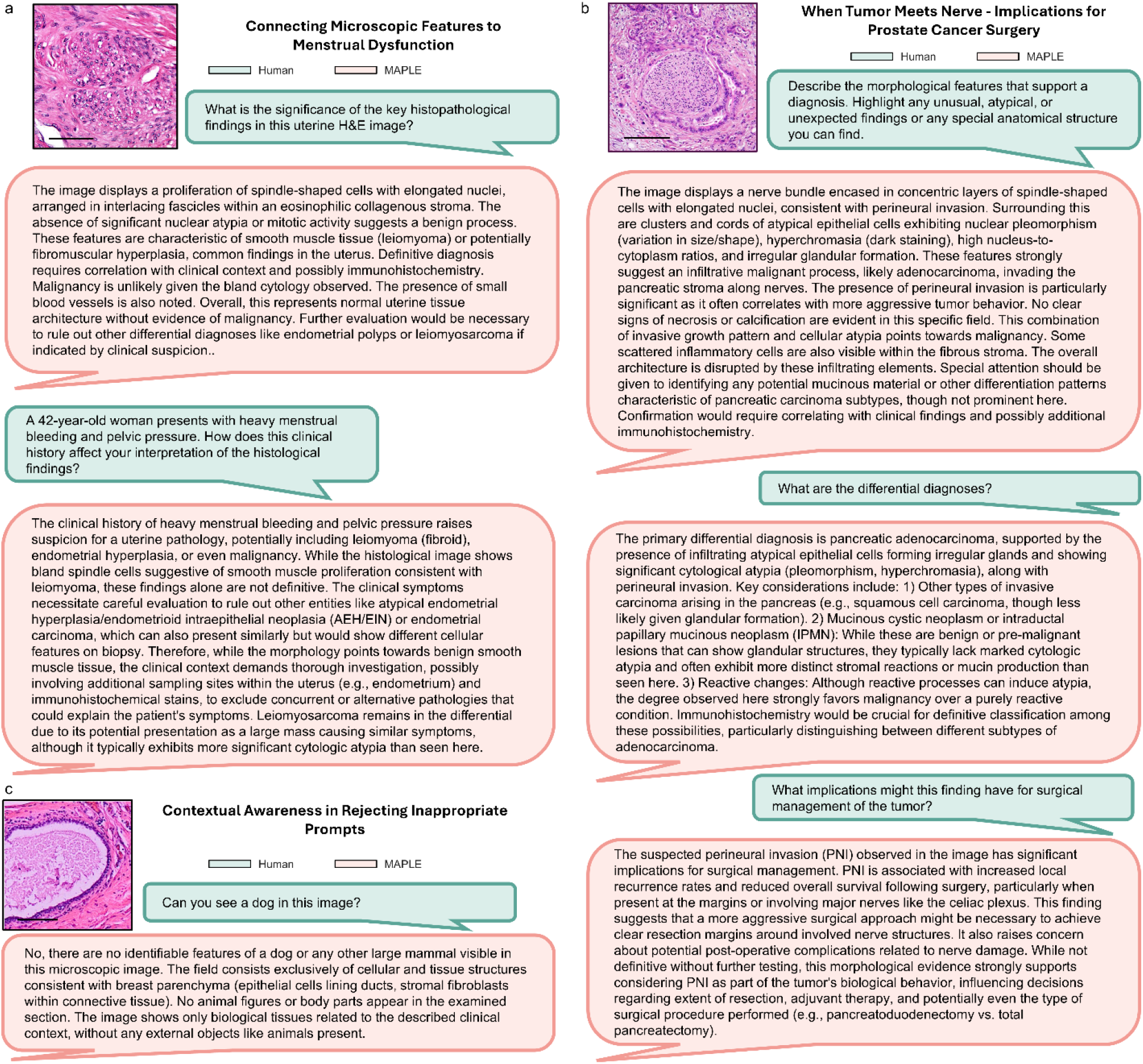
Context-aware clinical reasoning and interpretation of histopathological findings. **a,** Representative multi-turn dialogues illustrating MAPLE’s ability to relate uterine smooth muscle morphology to clinical symptoms. **b,** Recognize perineural invasion and discuss its surgical implications. **c,** Reject an unrelated visual query through model guardrails. User prompts are shown in green and MAPLE responses in pink.

**Supplemental Fig 3.**
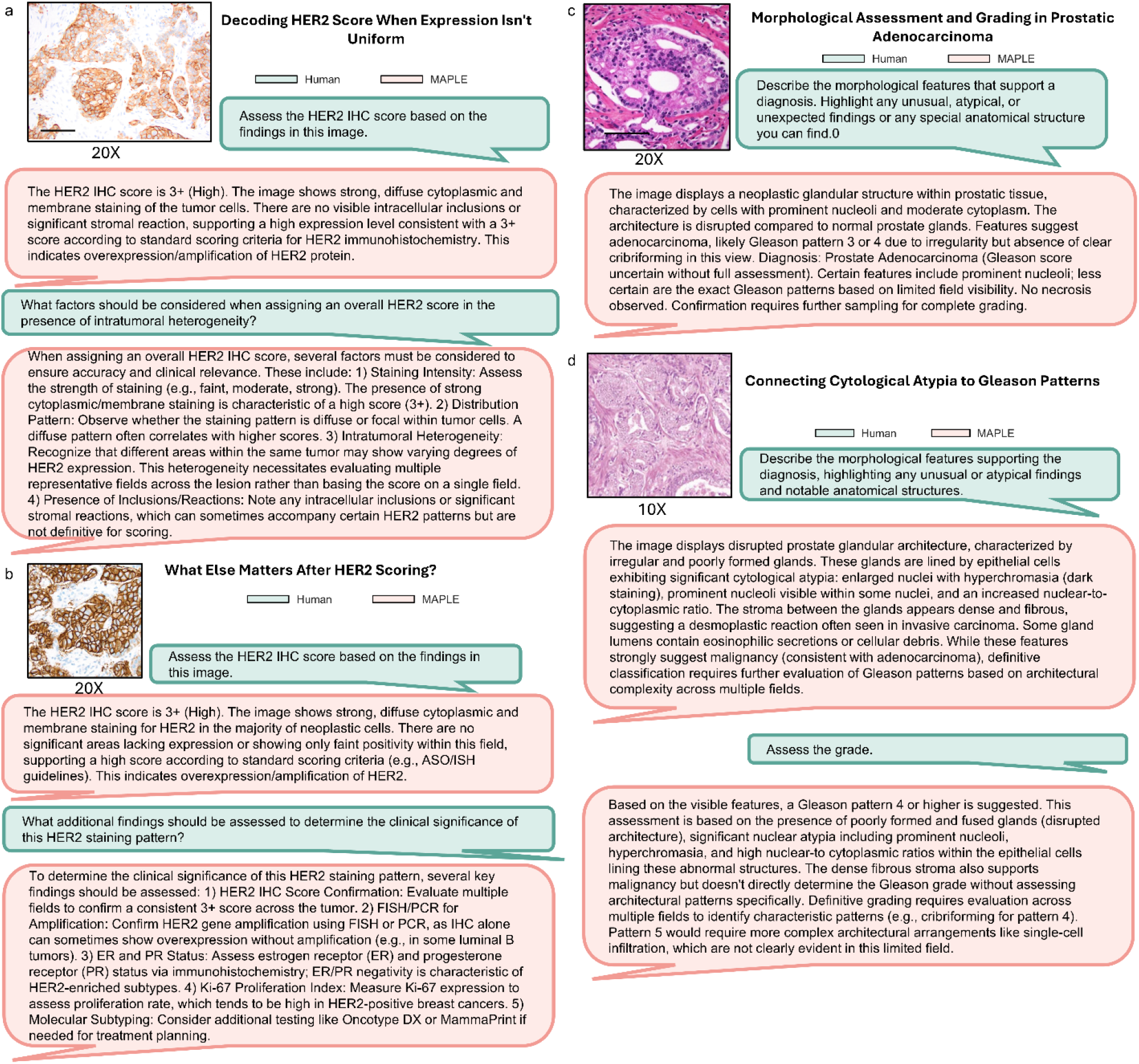
Interpretation of immunohistochemical findings and prostate cancer grading. **a,b,** Representative dialogues illustrating MAPLE’s assessment of human epidermal growth factor receptor 2 (HER2) immunohistochemical staining and additional diagnostic considerations. **c,d,** morphological assessment of prostatic adenocarcinoma, including nuclear atypia, glandular architecture, and Gleason grading. User prompts are shown in green and MAPLE responses in pink.

**Supplemental Fig 4.**
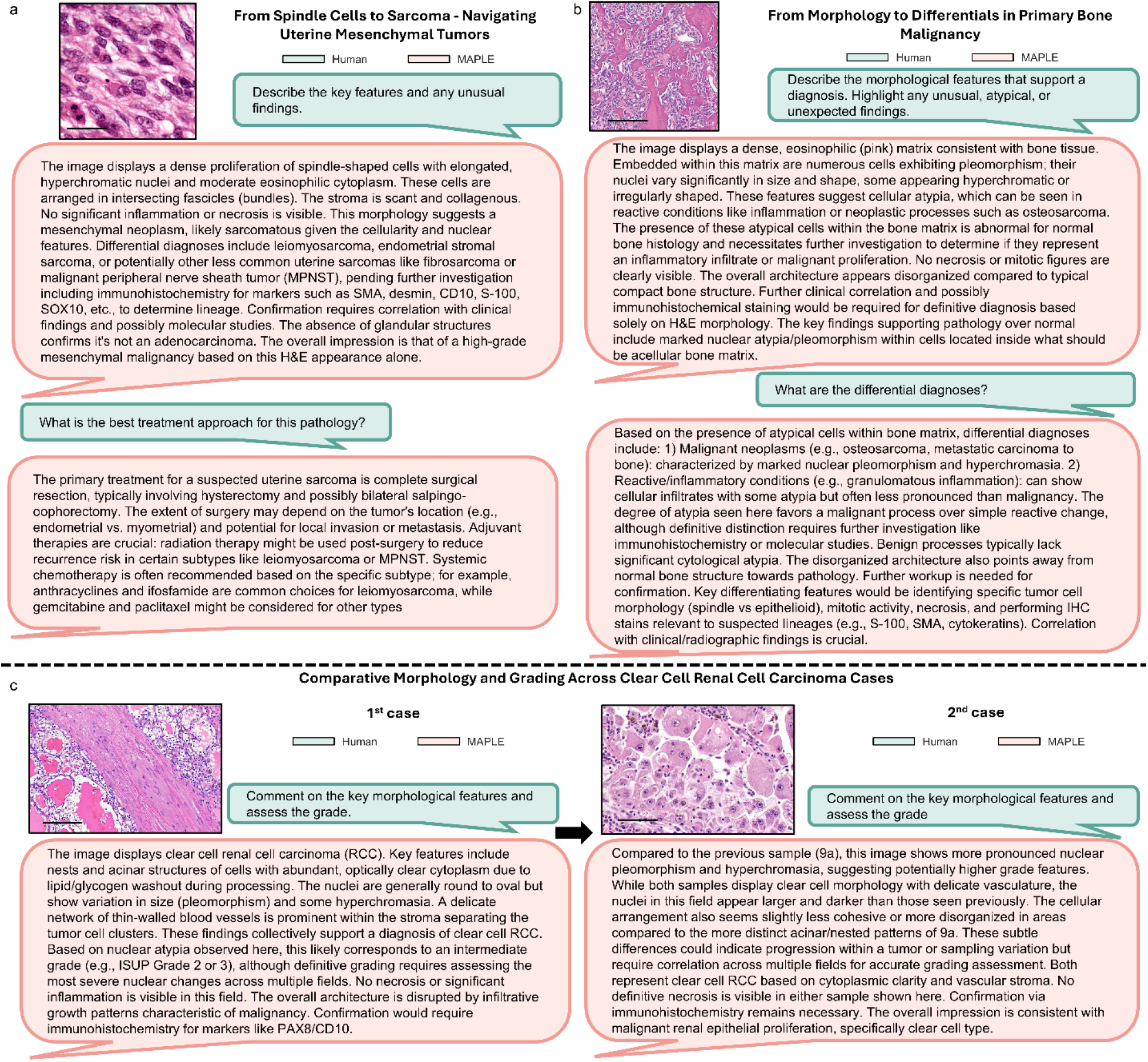
Diagnostic reasoning across mesenchymal tumors and renal cell carcinoma. **a,** Representative dialogues illustrating morphological assessment and treatment considerations for suspected uterine sarcoma. **b,** Differential diagnosis of a primary bone lesion. **c,** Comparative morphological evaluation and grading of two clear renal cell carcinoma cases (bottom). User prompts are shown in green and MAPLE responses in pink.

**Supplemental Fig 5.**
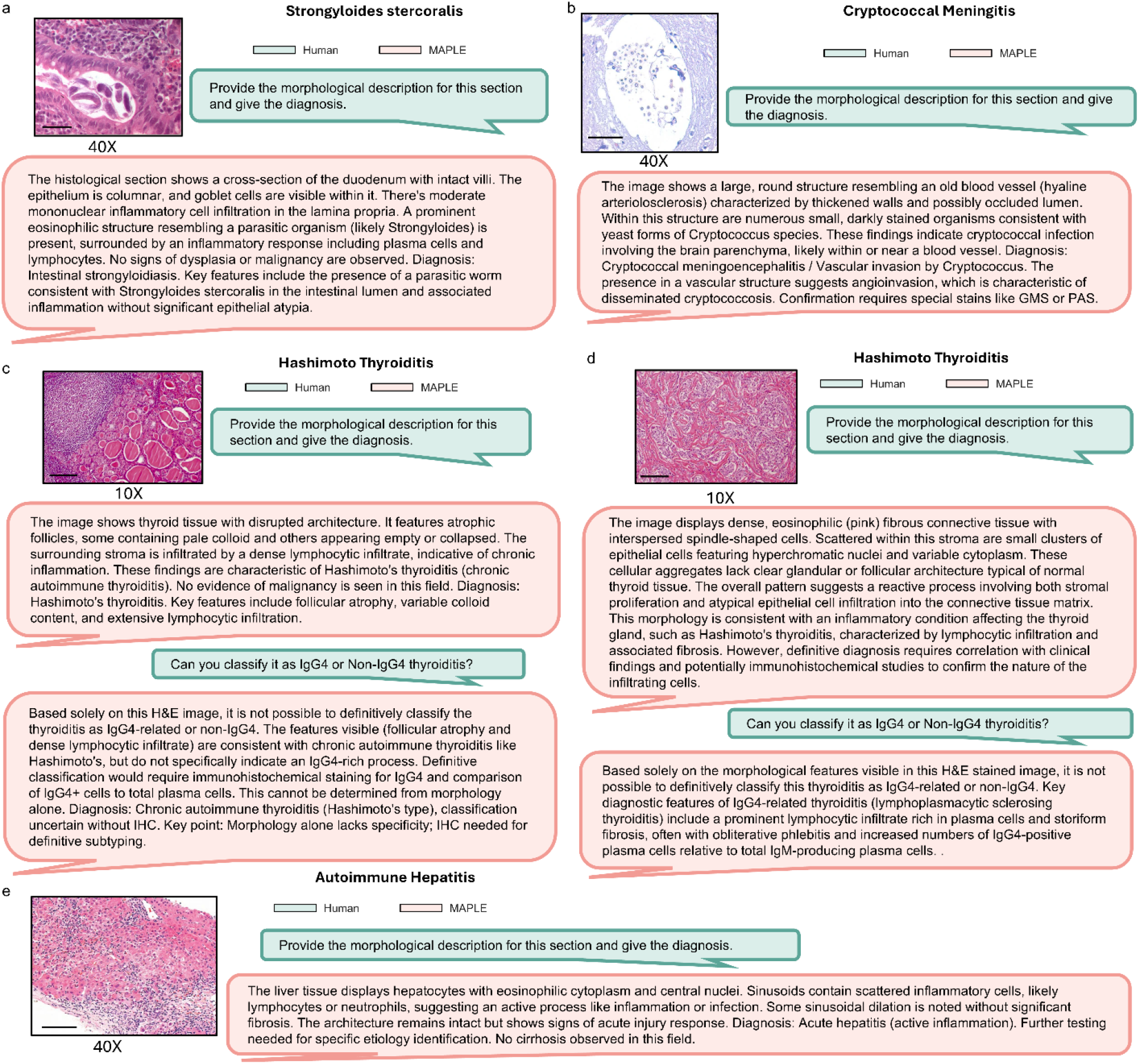
Diagnostic reasoning across infectious and autoimmune conditions. **a,** Representative dialogues illustrating morphological assessment and diagnosis of intestinal parasitic infection with *Strongyloides stercoralis*. **b,** Cryptococcal meningitis. **c,d,** Hashimoto’s thyroiditis with follow-up IgG4 subclassification queries. **d,** Autoimmune hepatitis. User prompts are shown in green and MAPLE responses in pink.

**Supplemental Fig 6.**
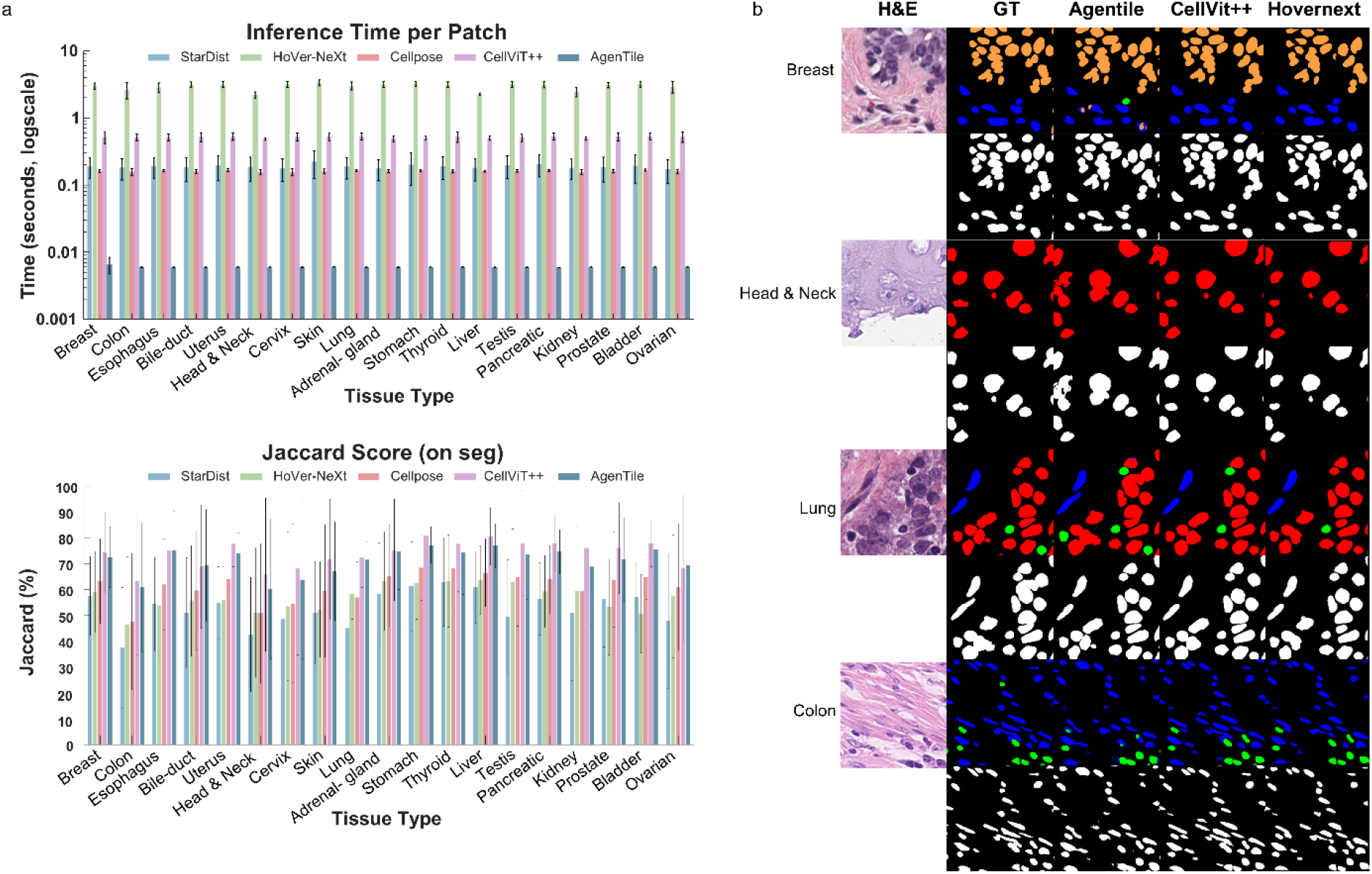
Tissue-specific performance and computational efficiency of AgenTile. **a,** Comparison of nuclei segmentation Jaccard scores (top) and inference times per patch (bottom, logarithmic scale) across 19 tissue types. **b,** Representative hematoxylin and eosin (H&E)-stained images from breast, head and neck, lung, and colon tissues, showing ground truth (GT) and predicted nuclei segmentation and classification results from AgenTile, CellViT++, and HoverNext.

## MATERIALS AND METHODS

### Curation of Maple Dataset

We curated a dataset for training MAPLE to respond to pathology-specific queries, comprising an alignment set of approximately 600,000 question-image-response pairs and an instruction-tuning set of approximately 100,000 images with multi-turn dialogues (2-4 question-answer pairs per image), both formatted following the LLaVA conversation structure. The dataset encompassed 10 organ systems, 25 tissue types, and 31 cancer subtypes.

Whole-slide images (N_WSI = 15,000) were downloaded from The Cancer Genome Atlas (TCGA) and manually curated into three quality categories: high-fidelity (N_WSI ≈ 6,500), backup (N_WSI ≈ 4,000), and low-fidelity (N_WSI ≈ 4,500). High-fidelity images exhibited consistent staining across the slide without processing artifacts, permitting patch extraction from any region. Backup images contained artifact-free regions suitable for targeted patch extraction should additional examples of certain tissue representations be required. Low-fidelity images, characterized by poor stain consistency, tissue folds, tears, air bubbles from cover slipping, or other processing artifacts, were excluded from training. TCGA data were used exclusively for training; no TCGA images were included in downstream evaluation or model testing.

Caption generation followed a multi-step process. For high-fidelity and backup images, existing captions were first obtained from the PathGen-1.6M dataset where available, which provided relevant *x,y* coordinates for patch extraction from whole-slide images. For images not represented in PathGen-1.6M, captions were generated by passing slide-level metadata through large language models (Qwen, Gemini), with corresponding patch regions determined through manual visual inspection. All captions, including those sourced from PathGen-1.6M, were further refined using Qwen and Gemini to ensure consistency, specificity, and removal of uninformative or overly generic descriptions. All data were de-identified before analysis.

### Curation of AgenTile dataset

AgenTile was trained on the PanNuke dataset, a semi-automatically generated dataset for nuclei instance segmentation and classification. PanNuke comprises 7,901 images (256x256 pixels) captured at 40x magnification (0.25 µm/pixel), containing 189,744 labeled nuclei with instance segmentation masks across 19 tissue types: adrenal gland, bile duct, bladder, breast, cervix, colon, esophagus, head and neck, kidney, liver, lung, ovarian, pancreatic, prostate, skin, stomach, testis, thyroid, and uterus. Each nucleus is annotated with one of five cell categories: neoplastic, inflammatory, connective, dead, and epithelial. The dataset exhibits class imbalance, with the dead nuclei category being particularly underrepresented. As patches were extracted from whole-slide images, some nuclei located at patch edges are cropped, with fewer than 10 visible pixels in certain cases.

The dataset is organized into three predefined folds. We used folds 1 and 2 for training (5,179 images) and held out fold 3 for benchmarking (2,722 images). Tissue representation varied across folds, with high representation for breast, colon, and esophagus, and limited representation for bladder, ovarian, and pancreatic tissues.

### Training of PathLLM

MAPLE integrates a pathology-specialized CLIP vision encoder with the Qwen3-14B language model through a two-layer MLP projector (PathLLM). The vision encoder (ViT-L/14) extracts patch features ***z***_*v*_ ∈ ℝ^1024^from the second-to-last transformer layer, where the subscript *v* denotes visual features. These features are projected to the language model’s embedding space via:

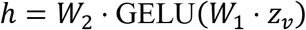

where *W*_1_ ∈ ℝ^5120×1024^and *W*_2_ ∈ ℝ^5120×5120^. are learned weight matrices, and the projected visual token ℎ ∈ ℝ^5120^. The projected visual features are concatenated with text token embeddings to form the multimodal input sequence:

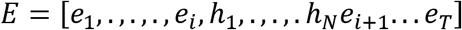

where *e*_*j*_denotes the embedding of the *j*-th text token (*j* = 1, …, *T*), *i* is the position at which visual tokens are inserted, *N* is the number of visual patch tokens, and *T* is the total number of text tokens.

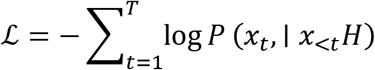

where *x*_*t*_ is the *t-*th text token, *x*_<*t*_ denotes all preceding tokens, and *H* denotes the full multimodal context comprising both visual and text tokens. Loss is computed only over text token positions.

Training followed a two-stage procedure. In Stage 1 (alignment pretraining), only the MLP projector was trained while the vision encoder and language model remained frozen. The model was trained on the alignment dataset (approximately 600,000 image-question-response pairs) for 1 epoch using the AdamW optimizer with learning rate 1e-3, cosine learning rate schedule, warmup ratio 0.03, and no weight decay. Training used a per-device batch size of 16 with gradient accumulation over 4 steps (effective batch size 64 per device), maximum sequence length of 2048 tokens, bfloat16 mixed precision, gradient checkpointing, and DeepSpeed ZeRO Stage 2 optimization.

In Stage 2 (instruction fine-tuning), the full model was trained on the instruction-tuning dataset (approximately 100,000 multi-turn dialogues) for 1 epoch. Training used learning rate 2e-5, cosine schedule, warmup ratio 0.03, per-device batch size of 2 with gradient accumulation over 16 steps (effective batch size 32 per device), maximum sequence length of 8192 tokens, bfloat16 precision, and DeepSpeed ZeRO Stage 3 optimization. Images were padded to preserve aspect ratio during fine-tuning. Training was performed on 8 NVIDIA A100 GPUs.

### Training of AgenTile

AgenTile adapts the CellViT architecture by replacing the original Vision Transformer backbone with EfficientSAM-Ti, a lightweight encoder optimized for segmentation tasks. The model processes 256x256 pixel patches at 40x magnification (0.25 µm/pixel) and produces four prediction outputs: binary nuclei segmentation, horizontal-vertical (HV) gradient maps for instance separation, nuclei type classification, and tissue type prediction.

The encoder employs EfficientSAM-Ti, consisting of 12 transformer blocks with embedding dimension 192, 3 attention heads, and MLP ratio of 4. Input images are divided into 16x16 pixel patches, yielding 16x16 spatial tokens.^61^ Intermediate features are extracted from transformer blocks 3, 6, 9, and 12 to provide multi-scale representations for skip connections. A two-layer neck projects the final encoder output to 256 dimensions. The decoder follows a U-Net-style architecture with shared skip connection processing and task-specific upsampling branches.^62^ Skip connections from the four extracted layers undergo progressive upsampling through transposed convolutions: the deepest features (layer 12) pass through a bottleneck upsampler, while shallower features (layers 9, 6, 3) are processed by dedicated decoder blocks before concatenation. Each task-specific branch (binary segmentation, HV map, nuclei type) contains its own upsampling path with Conv2D blocks (kernel size 3, with dropout) and transposed convolutions, progressively restoring spatial resolution from 16x16 to 256x256. The input image additionally passes through a shallow convolutional stem (3 to 32 to 64 channels) to preserve high-frequency details for the final skip connection. The binary segmentation branch outputs 2 channels (foreground/background). The HV map branch outputs 2 channels encoding horizontal and vertical gradients, which guide watershed-based instance separation during post-processing. The nuclei type branch outputs 5 channels corresponding to neoplastic, inflammatory, connective, dead, and epithelial cell categories. A linear classifier head predicts tissue type from mean-pooled encoder features.

We trained AgenTile for 130 epochs using AdamW optimizer with learning rate 3x10⁻⁴, weight decay 10⁻⁴, and batch size 64. Learning rate followed an exponential decay schedule (decay factor = 0.85). The encoder was frozen for the first 25 epochs, then unfrozen to allow end-to-end fine-tuning. We employed mixed-precision training (FP16) for computational efficiency. The total loss combines task-specific objectives:

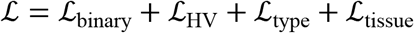

For binary segmentation, we used Focal Tversky loss combined with Dice loss:

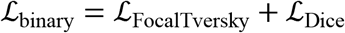

where Focal Tversky loss is defined as:

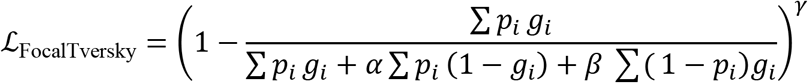

where the sum is taken over all pixels *i* in the patch, *p*_*i*_ ∈ [0,1] denotes the predicted foreground probability, *g*_*i*_ ∈ {0,1} denotes the binary ground truth label, *α* and *β* control the penalties for false positives and false negatives respectively, and *γ* is the focal exponent. HV map prediction employed mean squared error (MSE) and mean squared gradient error (MSGE):

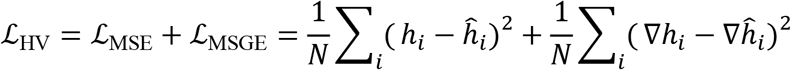

where the sum is taken over all pixels *i*, N_pix is the total number of pixels in the patch, ℎ_*i*_and ℎ^_*i*_ denote the ground truth and predicted HV map values respectively, and ∇ denotes the spatial gradient operator computed as finite differences. Nuclei type classification used binary cross-entropy, Dice loss, and multi-class Focal Tversky loss:

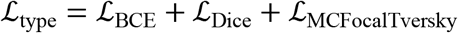

Tissue classification used cross-entropy loss:

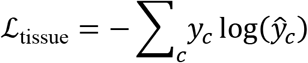

where the sum is taken over all C tissue classes, *y*_*c*_ ∈ {0,1} is the ground truth one-hot label for class *c*, and *y*^_*c*_ is the predicted probability for class *c*.

Training images underwent extensive augmentation including random rotation, horizontal and vertical flips, downscaling, Gaussian blur, additive noise, color jitter, superpixel perturbation, zoom blur, and elastic transformations. Instance segmentation maps were derived from model outputs using the HoVerNet post-processing pipeline, where binary predictions and HV maps were combined through marker-controlled watershed segmentation, with gradient information guiding the separation of touching nuclei. Each detected instance was assigned to a cell type based on the maximum probability from the nuclei type prediction map.

### Benchmarking image encoder embedding space

To evaluate the quality of learned visual representations, we compared MAPLE’s image encoder against state-of-the-art pathology foundation models (UNI2-h and Virchow2) on the BACH breast cancer classification dataset. The BACH dataset contains 400 histopathology images across four diagnostic categories: benign, in situ carcinoma, invasive carcinoma, and normal tissue. This evaluation assesses embedding quality independent of language model components, measuring how well each encoder captures diagnostically relevant visual features.

For each encoder, we extracted image embeddings and evaluated them using complementary protocols. Linear probe evaluation trained a logistic regression classifier on frozen embeddings using 5-fold stratified cross-validation, with embeddings standardized prior to training. We report accuracy, balanced accuracy, and macro F1-score. *k*-nearest neighbors (*k*-NN) classification assessed local neighborhood structure by evaluating classification accuracy at *k* ∈ {1, 3, 5, 10, 20} using cosine distance on L2-normalized embeddings.

Clustering quality metrics quantified class separability without supervised labels. Silhouette score measures how similar samples are to their own class compared to other classes, with higher values indicating better-defined clusters. The Calinski-Harabasz index computes the ratio of between-cluster to within-cluster dispersion, where higher values indicate more compact and well-separated clusters. The Davies-Bouldin index measures average similarity between clusters, with lower values indicating better separation. Retrieval metrics (Precision@*k*) evaluated whether nearest neighbors in embedding space share the same diagnostic class, computed by identifying the *k* most similar images for each query and calculating the proportion with matching labels.

We additionally computed intra-class and inter-class distance distributions using cosine distance. The separation ratio, defined as the ratio of mean inter-class to mean intra-class distance, quantifies how well the embedding space separates diagnostic categories:

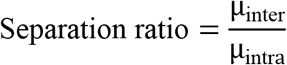

Where µ_inter_ and µ_intra_ denote the mean inter-class and intra-class cosine distances respectively, with higher values indicating better discrimination. Uniform Manifold Approximation and Projection (UMAP) visualizations provided qualitative assessment of cluster structure in two-dimensional projections.

### Benchmarking of image encoder on PathMMU

We evaluated MAPLE’s image encoder on PathMMU, a comprehensive pathology visual question answering benchmark comprising expert-curated multiple-choice questions across four domains: PubMed (scientific pathology documents), EduContent (educational YouTube teaching videos), SocialPath (Twitter posts by pathology experts), and Atlas (authoritative pathology textbooks and clinical guidelines). Each domain contains test sets with questions spanning diverse tissue types, staining methods, and diagnostic reasoning tasks.

For all CLIP-based models, we employed zero-shot visual question answering through image-text similarity matching. Given a pathology image and a multiple-choice question with four options (A, B, C, D), we encoded the image using the model’s image encoder and independently encoded each candidate answer by concatenating the question text with the option text. We computed cosine similarity between the normalized image embedding and each text embedding, selecting the option with maximum similarity as the predicted answer:

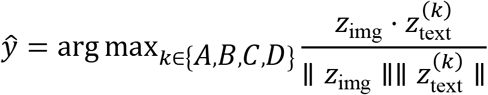

where *z*_img_denotes the image embedding and *z*^(*k*)^_text_denotes the text embedding for option *k*. This approach evaluates whether the learned visual representations capture semantically meaningful features that align with pathology-specific textual descriptions without task-specific fine-tuning.

We compared MAPLE against pathology-trained CLIP encoders (CONCH, PLIP, BiomedCLIP, MUSK, Quilt) and general-domain baselines (OpenAI CLIP ViT-B/16 and ViT-B/32). Accuracy was computed as the proportion of correctly answered questions within each domain, and we report performance stratified by domain to assess generalization across different pathology content sources.

### Benchmarking of PathLLM on PathMMU

We evaluated MAPLE on PathMMU, a pathology visual question answering benchmark comprising four domains: PubMed (scientific pathology documents), EduContent (educational YouTube teaching videos), SocialPath (Twitter posts by pathology experts), and Atlas (authoritative pathology textbooks and clinical guidelines). Each sample consists of a histopathology image paired with a multiple-choice question with four options (A–D).

For each question, we constructed the input prompt by concatenating the question text, answer options formatted on separate lines, and an instruction requesting structured output: "Please show your choice in the answer field with only the choice letter." The image token was prepended to the text following standard LLaVA formatting conventions, using the Qwen3 conversation template.

Generation used greedy decoding (temperature = 0) with repetition penalty 1.2, no-repeat *n*-gram size of 4, and maximum 4,096 new tokens.

Model outputs were parsed via regular expressions to extract predicted choice letters, matching single-letter responses (A, B, C, or D) at the output start, optionally followed by punctuation or newlines. Responses with multiple inconsistent matches or no valid match were counted as extraction errors. Accuracy was computed as correct predictions divided by total samples.

### Benchmarking of AgenTile and comparative models on PanNuke

We evaluated AgenTile and all comparative nuclei segmentation models on the PanNuke test set (fold 3; 2,722 images across 19 tissue types). For each 256x256 image patch at 40x magnification, models predicted binary nuclei masks, horizontal-vertical (HV) distance maps, and five-class nuclei type classification maps (neoplastic, inflammatory, connective, dead, epithelial). Instance segmentation was derived using HoVerNet post-processing with watershed segmentation guided by HV maps.

Predicted and ground truth nuclei instances were paired using centroid proximity matching with a radius of 12 pixels at 40x magnification, corresponding to approximately 3 μm. Paired instances constituted true positives (TP), unmatched ground truth instances were false negatives (FN), and unmatched predictions were false positives (FP). From these counts we computed instance-level detection metrics:

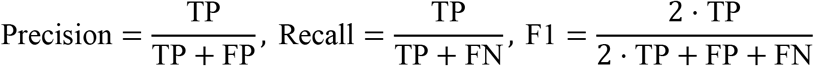

where TP, FP, and FN refer to instance-level detection counts based on centroid proximity matching. The instance-level Dice coefficient follows the same formulation as F1 at the instance level. The Jaccard index was computed as:

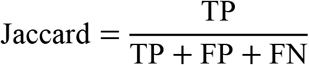

Given the clinical importance of identifying malignant cells, we separately evaluated neoplastic nucleus classification accuracy over successfully detected nuclei only. For each paired nucleus, the ground truth and predicted neoplastic versus non-neoplastic labels were compared; undetected ground truth nuclei were excluded from this metric. Classification accuracy was computed as:

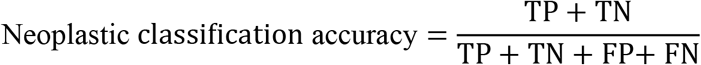

where TP, TN, FP, and FN here refer to classification outcomes among successfully paired nuclei only: TP denotes paired neoplastic nuclei correctly classified as neoplastic, TN denotes paired non-neoplastic nuclei correctly classified as non-neoplastic, FP denotes paired non-neoplastic nuclei misclassified as neoplastic, and FN denotes paired neoplastic nuclei misclassified as non-neoplastic. All metrics were stratified across the 19 tissue types in PanNuke, reporting mean ± standard deviation across constituent images for each tissue type.

### Benchmarking zero-shot classification

We evaluated the zero-shot classification performance of six vision-language foundation models: MAPLE, BiomedCLIP, CONCH, PLIP, QuiltNet, and MUSK. Each model enables classification via contrastive image-text alignment, such that histopathological images are assigned to the class whose textual prompt embedding maximizes cosine similarity within a shared representational space. Model performance was benchmarked across three publicly available histopathology datasets spanning heterogeneous tissue types and classification granularities: PatchCamelyon (327,680 lymph node patches with binary metastasis labels), SkinCancer (129,364 skin patches across 16 tissue and tumor classes), and BACH (400 breast histology microscopy images across four diagnostic classes). Because zero-shot inference involves no model training, all labeled images from each dataset were pooled for evaluation irrespective of official splits (for BACH, the 400 publicly labeled images were used). For each class, four to seven free-text prompts combining diagnostic terminology and morphological descriptors were encoded by the model’s text encoder; the resulting embeddings were L2-normalized, averaged, and re-normalized to yield a single class embedding, with identical prompt sets used across all six models:

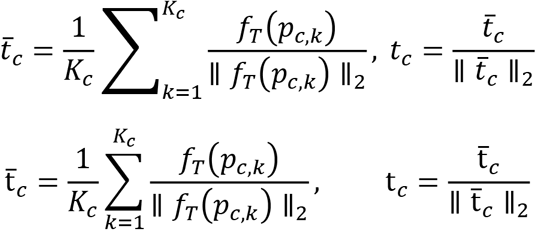

where *c* indexes the class, *K*_*c*_ is the number of prompts defined for class *c*, *f*_*T*_ denotes the model’s text encoder, *p*_*c*,*k*_ is the *k*-th prompt for class *c*, and t_*c*_ is the resulting normalized class embedding.

Text prompts for each class were defined based on pathological domain knowledge, describing tissue appearance, staining characteristics, and diagnostic terminology. Images were preprocessed with each model’s native pipeline, image embeddings were L2-normalized, and class probabilities were obtained by applying a softmax to the cosine similarities scaled by the model’s learned temperature; the predicted class was the argmax of these probabilities:

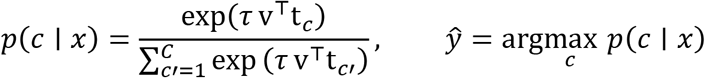

where *x* is the input image, v is its L2-normalized image embedding, *τ* is the model’s learned temperature parameter, *C* is the total number of classes, and *c*′ is a dummy index ranging over all classes in the normalising summation.

Performance was summarized by balanced accuracy and macro-averaged F1 score, with uncertainty estimated via patch-level nonparametric bootstrap (1,000 resamples with replacement):

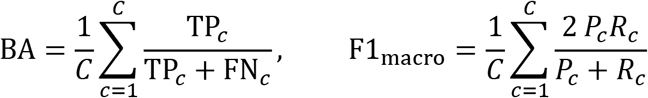

where TP_*c*_ and FN_*c*_ denote true positive and false negative counts for class *c* under a one-vs-rest scheme, and *P*_*c*_ and *R*_*c*_ denote the corresponding precision and recall for class *c*.

To assess performance on clinical material, the same six models were evaluated on 15 hematoxylin and eosin (H&E)-stained whole-slide images of breast tissue. An expert pathologist annotated regions of invasive carcinoma, ductal carcinoma in situ (DCIS), atypical lobular hyperplasia/lobular carcinoma in situ (ALH/LCIS), and benign tissue using Aperio ImageScope. Each slide was partitioned into a non-overlapping grid of 672x672-pixel patches at 40x magnification. To reduce computational overhead, tissue-containing patches were identified at the lowest-resolution pyramid level by pixel intensity thresholding: pixels with at least one RGB channel value below 220 were classified as tissue, and patches with less than 50% tissue coverage were discarded as white space. A patch was labeled positive (tumor) if its center fell within an invasive carcinoma annotation; patches whose centers fell within carcinoma in situ (DCIS or ALH/LCIS), benign tissue annotations, or unannotated tissue regions were labeled negative. When a patch center fell within overlapping annotations, priority was assigned in the order: invasive carcinoma > DCIS > ALH/LCIS > benign. This labeling scheme reflects the clinical objective of detecting invasive malignancy, with in situ lesions and benign tissue grouped together for binary classification.

Zero-shot inference followed the patch-level protocol described above, with seven prompts per class ensembled into a single embedding for each of the two classes (tumor and non-tumor). The softmax probability assigned to the tumor class served as the patch-level score. A single binary decision threshold per model was selected by maximizing Youden’s J statistic over all patches pooled across the 15 slides:

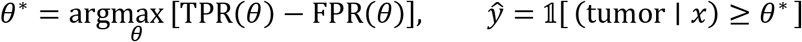

where *θ* is a candidate decision threshold, and TPR(*θ*) and FPR(*θ*) denote the true positive rate and false positive rate at threshold *θ*, respectively.

Balanced accuracy and macro-averaged F1 score were computed at this threshold. To assess generalization, we additionally performed leave-one-slide-out cross-validation, selecting the optimal threshold on 14 slides and evaluating on the held-out slide, repeating for all 15 slides. For spatial visualization, patch-level tumor probabilities were mapped back to their grid positions and overlaid on a thumbnail of each slide, and thresholded predictions were compared with the pathologist annotations to produce maps of true-positive, false-positive, false-negative, and true-negative patches.

### Patch selection of most diagnostically relevant patches

Each section was tiled at 40×, every nucleus was segmented and assigned to one of the five PanNuke classes, and the tiles were divided into 336 × 336-pixel candidate patches. A patch was eligible if neoplastic nuclei were its most frequent class, at least 90% of its area was tissue, it contained at least 20 nuclei, and the zero-shot tumor heat map assigned its location a tumor probability of at least 0.70. Eligible patches were ranked by the product of this probability and their neoplastic purity (the fraction of nuclei classified as neoplastic), and the 20 highest-ranked, spatially separated patches were retained. If fewer than 20 qualified, the most cellular of the remainder made up the number. The three retained patches with the highest tumor probability were passed to the language model together with an overview image of the section. Slides on which less than 1% of the tissue reached the probability threshold were treated as non-neoplastic and represented by their most cellular epithelial and stromal patches.

### AgenTile and image encoder ablation study

We assessed the precision with which AgenTile identified regions of interest. Each WSI was divided into a grid of non-overlapping 1008x1008-pixel patches at 40x magnification. AgenTile processed each patch, segmenting and classifying every nucleus into one of five types: neoplastic, epithelial, connective, inflammatory, or dead. Each processed patch was then subdivided into nine 336x336-pixel patches, and the number of nuclei of each type was counted within every patch. Patches were labeled based on their neoplastic cell content: those with greater than 90% neoplastic nuclei were labeled "neoplastic, high density," those with 70–90% neoplastic nuclei were labeled "neoplastic, medium density," and all remaining patches were labeled "benign." This density stratification, rather than a single binary label, was designed to provide the downstream language model with more granular context.

The neoplastic patches identified by AgenTile were compared against annotations from a board-certified pathologist. A patch was marked as a true positive (TP) if it fell within an annotated invasive carcinoma region, and as a false positive (FP) if it fell within a region annotated as benign, carcinoma in situ, or unannotated tissue. Precision was calculated as:

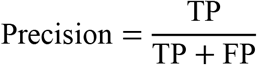

where TP and FP denote true positive and false positive patches as defined above. A weighted average precision was computed across the 15 breast tissue WSIs, with weights proportional to the number of neoplastic patches identified per slide, reflecting AgenTile’s overall ability to correctly prioritize regions of invasive malignancy for downstream processing by the language model.

## DATA AVAILABILITY

Data are available upon request from the corresponding author. The TCGA whole-slide images used for model training and curation are available at https://portal.gdc.cancer.gov/. The PathGen-1.6M dataset, from which existing captions and patch extraction coordinates were sourced for high-fidelity and backup images, is available at https://huggingface.co/datasets/jamessyx/PathGen. The PanNuke dataset used for nuclei instance segmentation and classification training is available at https://warwick.ac.uk/fac/cross_fac/tia/data/pannuke. The PathMMU benchmark dataset, encompassing the PubMed, SocialPath, EduContent, and Atlas evaluation domains, is available at https://huggingface.co/datasets/jamessyx/PathMMU. The BACH breast cancer histopathology dataset used for benchmarking is available at https://iciar2018-challenge.grand-challenge.org/. The PatchCamelyon dataset used for lymph node metastasis detection benchmarking is available at https://github.com/basveeling/pcam. The SkinCancer dataset used for 16-class dermatological classification benchmarking is available at https://github.com/openmedlab/Awesome-Medical-Dataset/blob/main/resources/SkinCancer.md. The 15 breast cancer whole-slide images used for in-house validation were acquired from the Johns Hopkins Department of Pathology and are not publicly available due to patient privacy constraints; access requests are subject to institutional review board (IRB) approval and data use agreement (DUA) requirements at Johns Hopkins University School of Medicine.

## CODE AVAILABILITY

The code used in this study will be made publicly available upon publication in a peer-reviewed journal.

## ACKNOWLEDGEMENTS

The authors acknowledge the following sources of support: UG3CA275681 (P.H.W.), UH3CA275681(P.H.W.), U54AR081774 (D.W.), U54CA268083 (D.W.). The contribution of A.M.B. was partially supported by the Spanish Ministerio de Ciencia, Innovación y Universidades and the Agencia Estatal de Investigación (MCIN/AEI/10.13039/501100011033), under grants PID2023-152631OB-I00 and AIA2025-164165-C41, co-financed by the European Regional Development Fund (ERDF), “A way of making Europe” and by the EU Horizon Europe research and innovation programme through the ESG4Stars project (GA n. 101290798)

## AUTHOR CONTRIBUTIONS

S.J., J.C., and D.W. conceived this project. S.J., J.C., R.Y. trained and developed the models. S.J., J.C., V.W., S.A.F., R.M., F.A.S., Y.S., A.F., T.D.G., D.K., E.A.H., P.N. conducted analysis. S.J., J.C., C.M., V.Q., K.L., V.D.A., D.N.S., I.W., F.W., A.F., V.M.R., L.D., R.A. downloaded, processed, or curated, data for training and/or benchmarking. R.H.H., A.M., A.L.K., and A.Y., provided technical feedback in the medical pathology space or computing space. D.W., A.M.B. and P.H.W provided ongoing scientific guidance and methodological input, critically reviewed the experimental approaches and results, and contributed to their interpretation. S.J., A.M.B., P.H.W and D.W. wrote the first draft of the manuscript, which all authors edited and approved.

## ETHICS STATEMENT

The authors declare no competing interests.

## Notes

### Competing Interest Statement

The authors have declared no competing interest.

